# Distinct alpha-synuclein strains derived from Parkinson’s disease patient tissues trigger differential inclusion pathology in a novel biosensor cell model

**DOI:** 10.1101/2025.04.01.646513

**Authors:** Anupam Raina, Wen Wang, Jose Carlos Gonzalez, Xiaohui Yan, Linda Overstreet-Wadiche, Jacques I. Wadiche, Chun-Li Zhang, Shu G. Chen

**Affiliations:** Department of Pathology, University of Alabama at Birmingham, Birmingham, Alabama, USA; Department of Molecular Biology, University of Texas Southwestern Medical Center, Dallas, Texas, USA; Department of Neurobiology, University of Alabama at Birmingham, Birmingham, Alabama, USA

**Keywords:** Parkinson’s disease, Dementia with Lewy bodies, Multiple system atrophy, Alpha-synuclein, RT-QuIC, Biosensor cells, Seeding, NEUROD1, Skin

## Abstract

**Background:** α-Synuclein (αSyn) can misfold and aggregate to form fibrillar ß-sheet-rich aggregates (“strains”) that are phosphorylated (p-αSyn) and deposited into intracellular inclusions in the brain, the pathological hallmark of synucleinopathies including Parkinson’s disease (PD), dementia with Lewy bodies (DLB), and multiple system atrophy (MSA). Previously, we reported that seed amplification assays such as real-time quaking-induced conversion (RT-QuIC) amplifies and detects αSyn strains from the patient skin. However, whether skin-derived αSyn strains induce disease-specific pathological features in a biological system is unknown.

**Methods:** We generated a U251 human glioblastoma cell line expressing fluorescently tagged αSyn carrying the PD-linked A53T mutation and Förster resonance energy transfer (FRET)-based U251 biosensor cells. Using fluorescence microscopy coupled with *in situ* detergent extraction, FRET-Flow cytometry and high-content confocal imaging, we examined the pathological burden and morphology of p-αSyn inclusions seeded by RT-QuIC-amplified patient skin and brain αSyn strains in αSyn-expressing U251 cells, FRET-based αSyn biosensor cells and αSyn biosensor cell-derived neurons.

**Results:** U251 cells allow robust and rapid *in situ* detection of detergent-insoluble intracellular αSyn inclusions triggered by exogenous αSyn seeds. In U251 FRET-based biosensor cells, PD skin-amplified strains induce a greater pathological burden and distinct p-αSyn inclusion morphology from PD brain-amplified and DLB skin-amplified strains. Inclusion morphology of DLB and MSA skin- and brain-amplified strains are comparable. Furthermore, skin-amplified αSyn strains induce neuronal inclusions and cause degeneration of induced neurons reprogrammed from the U251 biosensor cells. Finally, biosensor cell-propagated PD skin αSyn strains induce higher seeding activity measured by RT-QuIC than PD brain and DLB skin αSyn strains, linking intracellular pathological burden to *in vitro* seeding activity.

**Conclusions:** We report the detection of distinct PD strains derived from patient skin and brain tissues that trigger unique pathological phenotypes in U251 αSyn biosensor cells and cause degeneration of reprogrammed neurons from the same cell lineage. Moreover, DLB and MSA skin αSyn strains mimic pathological features of their brain αSyn strains in these biosensor cells. Therefore, the U251 αSyn biosensor cell model is a robust tool to potentially discriminate PD and DLB synucleinopathies and to study αSyn tissue- and strain-specific etiology and pathogenesis.

**Graphical abstract:** 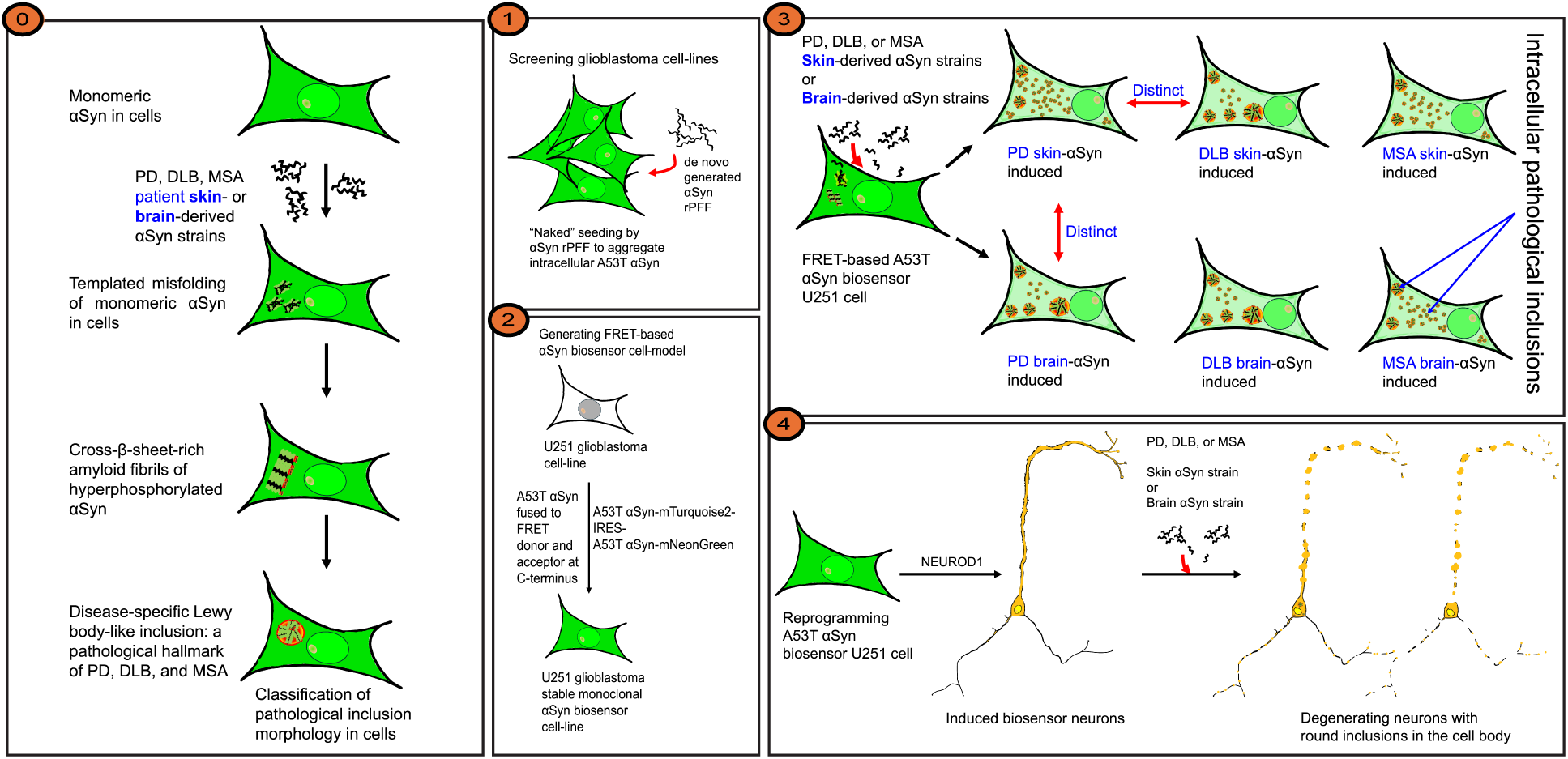

## Background

The accumulation of misfolded and aggregated alpha-synuclein (αSyn) in the brain is a pathological hallmark of a group of neurodegenerative diseases collectively called synucleinopathies, which include Parkinson’s disease (PD), dementia with Lewy bodies (DLB), and multiple system atrophy (MSA) [1–5]. Aggregated αSyn is invariably present in neuronal inclusions known as Lewy bodies and Lewy neurites in PD and DLB [1], and in glial cytoplasmic inclusions known as Papp-Lantos bodies in MSA [2–4]. Lewy bodies and Lewy neurites are round, oval or irregularly shaped filamentous inclusions present in the cytosol of neurons in the substantia nigra, cingulate cortex, and other brain regions [1]. In MSA, glial cytoplasmic inclusions are primarily found in the cytosol of oligodendrocytes, whereas filamentous inclusions are occasionally detected in neurons with round, oval, C-shape or ring-like inclusions in the frontal cortex, dentate fascia, substantia nigra, or pontine nucleus [2–4]. More than 90% of aggregated αSyn in these pathological inclusions is phosphorylated at serine 129 (p-αSyn) in a diseased brain, whereas in a healthy brain only about 4% of αSyn is phosphorylated at this site [6,7]. Studying cellular mechanisms underlying the formation of distinct p-αSyn-containing intracellular inclusions may further our understanding of pathogenesis of individual synucleinopathies.

αSyn is an intrinsically disordered protein [8] that can adopt different conformations [9,10]. It is enriched and localized in the presynaptic terminals of neurons and is presumably involved in the recycling of synaptic vesicles and synaptic transmission [11,12]. In a healthy intracellular environment, soluble αSyn exists as a natively unfolded monomer [13]. However, in a pathological intracellular environment, monomeric soluble αSyn tends to aggregate into toxic cross-ß-sheet-rich oligomers and assemblies [14–18] that is a characteristic of an amyloid [19,20].

Aggregated αSyn assemblies have been shown to behave as a strain with prion-like properties, which was first shown to propagate from brain into grafted neurons in PD patients [21,22], followed by neuron-to-neuron or cell-to-cell [23,24] propagation, and transmission between different brain regions *in vivo* [25,26]. Exogenous αSyn strains can also seed the aggregation of intracellular αSyn into Lewy body-like inclusions in cultured cells and neurons [26–31].

In prior studies, *de novo* generated cross-ß-sheet-rich fibrils using recombinant αSyn [32,33] (commonly known as recombinant pre-formed fibrils or rPFF) have been used by investigators to study PD pathogenesis in cultured cells and neurons [27,28]. rPFF can exist as polymorphic assemblies that possess different toxicity and seeding activity *in vitro*, in cells, in neurons and *in vivo* [34–36]. These studies led to the hypothesis that distinct strains of αSyn might underlie the pathological and clinical heterogeneity observed in synucleinopathies. Consequently, rPFF, patient brain homogenate or αSyn assemblies amplified from patient brain homogenate by protein misfolding cyclic amplification assay (PMCA) (another seed amplification platform) were used to study disease-specific pathology, biochemical characterization of conformers by partial digestion with proteinase-K, seeding activity *in vitro*, *in vivo*, and in cells or neurons [37–42]. These studies in combination with cryo-electron microscopy (cryo-EM) studies revealed that distinct pathologies were triggered by brain homogenates and brain-amplified αSyn strains in comparison to rPFF [43–45], suggesting the importance of using patient-derived αSyn strains for studying disease pathology. Both patient brain homogenate and brain homogenate-amplified αSyn strains have been studied. Using PMCA, brain-amplified PD and MSA strains induce higher p-αSyn pathology than brain-amplified DLB strains whereas MSA brain homogenate induces higher p-αSyn pathology than PD and DLB brain homogenate *in vivo* [40], confirming the existence of different αSyn strains in PD, DLB, and MSA. Surprisingly, PD patient brain homogenate seeds intracellular A53T αSyn differentially in the same cell-model as reported by two independent studies using the human embryonic kidney cell line (HEK 293T) [37,38]. Given that intracellular milieu can affect pathology of αSyn strains [46,47], it is reasonable to consider that the embryonic kidney cells might not be an ideal cell-model to study brain associated diseases like PD, DLB, and MSA. Taken together, these findings suggest that more than one αSyn strains are involved in the etiology and pathogenesis of PD, DLB, and MSA, and it is imperative to develop a cell model that closely mimics αSyn strain-specific pathology.

We have previously reported that pathological αSyn aggregates in PD, DLB, and MSA patient skin can be robustly detected by RT-QuIC assay of patient skin homogenates *in vitro* [48–50]. However, disease-specific pathological features triggered by skin αSyn strains in a biological system remain unknown.

In this study, we first aimed to generate a human brain-derived cell model that closely mimics αSyn inclusion pathology under physiological conditions. We screened different human glioblastoma cell lines that rapidly forms detergent-insoluble intracellular p-αSyn inclusions upon seeding by exogenous rPFF and generated a Förster resonance energy transfer (FRET)-based stable monoclonal αSyn biosensor U251 glioblastoma cell line, which was also amenable to reprogramming toward functional neurons by Neuronal Differentiation 1 (NEUROD1) transcription factor. Then, we investigated the biological and pathological features of skin-amplified and brain-amplified αSyn strains from PD, DLB, and MSA patients in U251 biosensor cells and U251 biosensor cell-derived neurons using FRET-Flow cytometry, high-content confocal imaging, and RT-QuIC assay. The morphology of intracellular p-αSyn inclusions was meticulously classified in U251 biosensor cells. We report that PD patient skin-amplified αSyn strains trigger higher intracellular pathological burden of p-αSyn inclusion and distinct morphology in comparison to the PD patient brain-amplified and DLB patient skin-amplified αSyn strains. Inclusion morphology of DLB and MSA patient skin-amplified αSyn strains remain conserved and mimic those of DLB and MSA patient brain-amplified αSyn strains. We also find that patient skin-and brain-amplified αSyn strains are capable of inducing neuronal inclusions and cause degeneration in U251 reprogrammed neurons. Following propagation of patient skin-derived αSyn strains in U251 biosensor cells and subsequent isolation of intracellular αSyn aggregates, we find that biosensor cell-derived PD skin αSyn strains induce higher seeding activity *in vitro* than DLB skin αSyn strains, suggesting a strong correlation between strain-specific intracellular pathological burden and its seeding activity. We conclude that the U251 FRET-based αSyn biosensor cell model is a robust tool to study disease-specific etiology and pathogenesis of synucleinopathies driven by αSyn strains derived from patient tissues.

## Methods

### Human tissue samples

Frozen tissue samples were obtained the NIH NeuroBiobankfrom neuropathologically confirmed cases of PD (n = 2 from scalp skin; n = 1 from brain cortex), DLB (n = 2 from scalp skin; n = 1 from brain cortex), MSA (n = 2 from scalp skin; n = 1 from brain cortex), and non-neurodegenerative (healthy) controls (n = 1 from scalp skin; n = 1 from brain cortex). Informed consent and Institutional Review Boards (IRB) approval for tissue collection and distribution were obtained by NeuroBioBank. As these tissue samples were de-identified and from deceased individuals, no further ethical approval was required.

### Tissue homogenization

Skin and brain tissue homogenates (10% w/v) were prepared at 4°C as described previously [50]. Briefly, skin tissue was first thawed and then rinsed three times in ice-cold 1X Dulbecco’s Phosphate-buffered saline (D-PBS) (Gibco) in 1.5ml tubes until blood was no longer visible, followed by mincing with a surgical blade into small pieces. Brain tissue was used directly. Both tissues were then homogenized at 1:10 (w/v) in the homogenization buffer containing 1X D-PBS, 150mM NaCl (Invitrogen), 1% Triton X-100 (BioRad), 5mM EDTA pH 8.0 (0.5M stock, Invitrogen), and 1X complete mini EDTA-free protease inhibitor cocktail (Roche Diagnostics) with 0.7 mm zirconium beads (BioSpec Products) in a mini-Beadbeater-16 device (BioSpec Products, Bartlesville, OK, USA) at 4°C for 5 cycles with 1 minute on and 3 minutes off. Homogenates were then centrifuged at 500xg for 5 minutes at 4°C. Supernatant was collected as clarified homogenate and a 10-fold (skin homogenate) and 1,000-fold (brain homogenate) dilutions were prepared in a solution containing 1X N2 (100X stock, Gibco) in D-PBS (N2/PBS). Clarified homogenate and diluted samples were immediately stored at -80°C until further use for RT-QuIC assay.

### Amplification of αSyn strains from patient tissue homogenates by RT-QuIC

αSyn strains from skin or brain homogenates were amplified by real-time quaking-induced conversion (RT-QuIC) assay. Reagents for RT-QuIC assay include 0.5M sodium phosphate pH 8.0 (J60825.AK, Thermo) stored at room temperature, 5M NaCl (AM9759, Invitrogen) stored at 4°C, thioflavin T (ThT) (596200, Sigma) stored at -20°C, and HPLC-grade water (W5-4, Fisher) stored at 4°C. RT-QuIC assay was adapted and performed as previously described [50]. Each well of black 96-well plate with flat optical bottom (265301, Thermo) was loaded with six 800 µm low-binding silica beads (BLBG-800-200-03, OPS Diagnostics) in a clean benchtop workstation. 1mg of lyophilized recombinant human αSyn (S-1001-2, lot. 051722AS, rPeptide,) was reconstituted at 1mg/ml and filtered through 0.5 ml Amicon 100 kDa filter (UFC510096, Millipore) at 10,000xg for 10 minutes at 4°C to get rid of aggregated αSyn, if any. RT-QuIC reaction mix containing 40 mM sodium phosphate, pH 8.0, 170 mM NaCl, 20 µM ThT in HPLC-grade water was prepared from the stock solutions as specified above and filtered through Millex-GV 0.22 µm PVDF membrane syringe filter (SLGV004SL, Millipore) into a sterile 25 ml boat while using benchtop air ionizer (963E, Static Control Solutions) to remove static charges from the work bench. Filtered recombinant αSyn was applied to the reaction mix so that the final concentration of αSyn in the reaction mix is 0.1 mg/ml. 98 µL of the reaction mix and 2 µL of the diluted skin or brain homogenate was applied to each well of 96-well plate containing silica beads. The plate was sealed with Nunc clear sealing film (Thermo) and loaded into BMG FLUOstar Omega plate reader (BMG Labtech). The RT-QuIC reaction was run at 42°C with 1 minute double-orbital shaking at 400 rpm and 1 minute rest, and optical gain was set at 1800. ThT fluorescence (448/10 nm excitation and 482/10 nm emission) was recorded every 45 minutes. RT-QuIC run was stopped when ThT fluorescence of healthy control samples reached a cut-off value of 50,000 relative fluorescence units (RFU). Each sample was run in triplicates and was considered positive if at least 2 out of 3 exceeded the cut-off RFU. Average ThT fluorescence was calculated for each sample regardless of whether they exceed the cut-off RFU. Raw data of ThT fluorescence was reported for each sample. To generate skin-amplified and brain-amplified αSyn fibrils, ThT was not used in the RT-QuIC reaction mix, however, amplification was simultaneously monitored in three separate wells with ThT in the reaction mix. After the RT-QuIC reaction was stopped, wells that did not contain ThT in RT-QuIC reaction mix were pooled in a 1.5 ml tube (357448, Beckman Coulter). Tubes were placed in a fixed-angle rotor (TLA-55, Beckman Coulter) and spun at 186,000xg for 30 minutes in an ultracentrifuge (393315, Beckman Coulter). Pellet was carefully rinsed with 1 ml of cold D-PBS, and spun again at 186,000xg for 30 minutes. The pellet was suspended in 20µL of cold D-PBS, and vortexed. Purified skin- and brain-amplified fibrils were stored in 1.5 ml Protein LoBind tubes (022431081, Eppendorf) at 4°C until further use. An aliquot of 3.5 µL of these skin- or brain-amplified αSyn fibrils were diluted in a final volume of 15 µL of 1X N2/PBS in 1.5 ml polystyrene tubes (53072, Active Motif), which was placed in a water bath pre-set to 4°C in a cup-horn sonicator (Q700, 431C2, QSonica). 10 minutes of sonication was set and carried out at an amplitude of 70 with 30 seconds of sonication and 30 seconds of rest in each cycle with 250-350 W power output per cycle. Sonicated αSyn fibrils were briefly spun down, and 1X N2/PBS was applied to each tube to make a final volume of 35 µL. After vortexing, 2 µL of sonicated samples were applied to 98 µL of RT-QuIC reaction mix in each well with or without ThT in a 96-well plate, followed by amplification and purification as described above. These patient-derived and amplified αSyn fibrils were used in cell culture experiments. To detect *in vitro* seeding activity of αSyn strains from the cell culture supernatant, the supernatant was diluted 1000 times in 1X N2/PBS. 2 µL of diluted supernatant was then applied to 98 µL of RT-QuIC reaction mix followed by RT-QuIC as described above. To detect *in vitro* seeding activity of αSyn strains from the sarkosyl-insoluble fraction of cells, 2 µL of the purified fraction was directly applied to 98 µL of RT-QuIC reaction mix followed by RT-QuIC as described above.

### Generation of recombinant αSyn preformed fibrils (rPFF)

For *de novo* generation of αSyn rPFF, 1 mg of lyophilized human recombinant αSyn was reconstituted in 500 µL of HPLC water in 1.5 ml Protein LoBind tube and agitated at 1,000 rpm for 7 days at 37°C in a thermomixer (EP5384000012, Eppendorf). The *de novo* generated rPFFs were purified using an ultracentrifuge as described above and resuspended in 400 µL of D-PBS and stored at 4°C until further use.

### Protein concentration

Concentration of purified skin and brain-amplified fibrils, rPFF, or cell lysates were measured by bicinchoninic acid (BCA) assay using bovine serum albumin as a standard. Additionally, absorbance at 205 nm with the SCOPES method (A_205_) was also used to measure αSyn fibril concentration on Nanodrop (ND-ONEC-W, Thermo Fisher) from 1.5 µL of sample and using 1.5 µL of D-PBS as a blank. Samples were vortexed for 5-10 seconds just before measuring concentrations in triplicates on Nanodrop. The mean concentration from triplicates was considered as final concentration of fibrils in that sample. Protein concentrations measured using BCA assay and Nanodrop A_205_ were found to be comparable, henceforth, Nanodrop A_205_ was used to measure fibril concentration in each sample.

### Transmission Electron Microscopy (TEM)

For negative staining to image skin or brain-amplified αSyn fibrils or rPFF, first, samples were diluted in HPLC water to a final concentration of 0.2 µg/µl. Formvar/carbon coated 200 mesh copper grid (FCF200-Cu-50, Electron Microscopy Sciences-EMS) was placed on PELCO TEM grid-holder (Ted Pella) and glow discharged for 25 seconds at 20 mA using PELCO easiGlow discharge apparatus (Ted Pella). After fixing the grid in a tweezer, 5 µL of diluted sample was applied on the grid and incubated for 1 minute. Excess sample was wicked off from the bottom of the grid using torn edge of Whatman filter paper. 7.5 µL of 1% uranyl acetate was applied on the grid and incubated for 45 seconds. Excess solution was wicked off from the edge of the grid using torn edge of Whatman filter paper. Then, grid was incubated in HPLC water drop for 30 seconds and excess water was wicked off. Grid was examined using JEOL 1400 FLASH TEM at 120 kV (JEOL USA Inc.) and digital images were captured using an AMT NanoSprint43L Mark-II camera (AMT Imaging).

For staining, sectioning, and imaging intracellular ultrastructure, first, U251 glioblastoma cells expressing A53T αSyn-EGFP were spun down at 400xg for 3 minutes. The cell pellet was suspended in fixing solution containing 2% glutaraldehyde in 0.15 M sodium cacodylate buffer (pH 7.4) for at least 2 hours at 4°C followed by rinsing twice for 15 minutes each in 0.15 M sodium cacodylate buffer. Cell pellet was then treated with 1% osmium tetroxide (EMS) in 0.15 M sodium cacodylate buffer in dark at room temperature for 1 hour followed by rinsing thrice for 15 minutes each. Pellet was then incubated in 1% low molecular weight tannic acid (Ted Pella) followed by rinsing thrice for 15 minutes each. Cell pellet was the dehydrated through graded 50%, 80%, 95%, and 100% ethyl alcohol treatments and transitioned through 100% propylene oxide to 50% propylene oxide in embedding resin to 100% embedding resin. Cell blocks were polymerized overnight at 60°C. Using silver to pale gold color interference, 70-90 nm thin sections were cut from cell blocks using a diamond knife (Diatome, EMS) on an ultramicrotome (Leica EM UC7). After drying, sections were mounted on 200 mesh copper grids and stained with 1% uranyl acetate and Reynold’s lead citrate for contrast. Grids were examined using JEOL 1400 FLASH TEM at 120 kV and digital images were captured using an AMT NanoSprint43L Mark-II camera.

### Plasmids

Lentiviral vectors used in this study were constructed using the plasmids obtained from Addgene. Constitutive expression of transgenes in glioblastoma cells using lentiviral vectors was driven by CMV promoter. The original plasmids used in this study were as following: pMDLg/pRRE (Addgene plasmid # 12251), pMD2.G (Addgene plasmid # 12259), and pRSV-Rev (Addgene plasmid # 12253) were a gift from Didier Trono [51]. pLV-eGFP (Addgene plasmid # 36083) was a gift from Pantelis Tsoulfas [52]. EGFP-alphasynuclein-A53T (Addgene plasmid # 40823) was a gift from David Rubinsztein [53]. pLenti-dsRed_IRES_SNCA:EGFP (Addgene plasmid # 92195) was a gift from Huda Zoghbi [54]. mTurquoise2-N1 (Addgene plasmid # 54843) was a gift from Michael Davidson and Dorus Gadella [55]. ER-mNeonGreen (Addgene plasmid # 137804) was a gift from Dorus Gadella [56]. EF1a_NEUROD1_P2A_Hygro_Barcode (Addgene plasmid # 120466) was a gift from Prashant Mali [57]. Lentiviral vectors that were constructed are as following: CMV-A53T αSyn-EGFP-WPRE-bGH poly(A), CMV-WT αSyn-EGFP-WPRE-bGH poly(A), CMV-A53T αSyn-mTurquoise2-IRES-A53T αSyn-mNeonGreen-PGK-Z418-WPRE-bGH poly(A), CMV-A53T αSyn-mTurquoise2-IRES-PGK-Z418-WPRE-bGH poly(A), CMV-IRES-A53T αSyn-mNeonGreen-PGK-Z418-WPRE-bGH poly(A), CMV-dsRed-IRES-NEUROD1-PGK-Z418-WPRE-bGH poly(A), and CMV-dsRed-IRES-PGK-Z418-WPRE-bGH poly(A) where EGFP is enhanced green fluorescent protein, WPRE is woodchuck hepatitis virus posttranscriptional regulatory element, bGH poly (A) is bovine growth hormone polyadenylation signal. All constructed lentiviral vectors were sequenced to confirm the integrity of transgenes.

### Cell culture

Human glioblastoma cell lines U251 (9063001, Sigma), U118, U87, U138, and DBTRG-05, as well as human embryonic kidney cell line Lenti-X HEK293T (632180, Takara) were maintained in 10 ml of growth medium consisting of DMEM (SH30243.01, Cytiva) with 10% fetal bovine serum (FBS) (Corning) and 1% penicillin/streptomycin (P/S) (15140122, Gibco) in a 10 cm dish at 37°C in a humidified incubator. Cells were passaged twice a week. For passaging, cells were rinsed once with D-PBS and dissociated from the dish by incubating with 1 ml of 0.05% trypsin (25-052-CV, Corning) for 3 minutes at 37°C. Trypsin was deactivated with 7 ml of growth medium applied to the cells and centrifuged at 400xg for 3 minutes at room temperature. Supernatant was discarded and pellet was suspended in required volume of medium for passaging in a fresh 10 cm dish with 10 ml of growth medium.

### Lentivirus production and purification

Third generation lentivirus was produced using Lenti-X HEK293T cells. Expanding Lenti-X 293T cells for lentivirus production was in part previously described [58,59]. First, cells were grown up to 60-70% confluency (5-8 million cells) in a 10 cm dish in growth medium. Medium was changed to 7 ml of DMEM medium, and cells were transfected with lentiviral transfer plasmid (4 µg) along with two packaging plasmids (pMDLg/pRRE: 2 µg, pRSV-REV: 0.5 µg), one envelope plasmid (pMD2.G/VSV-G: 1 µg), and 22.5 µL of 1 µg/µl of polyethylenimine in 1 ml of DMEM medium. Ratio of plasmid to polyethylenimine used was 1 to 3. Transfection was carried out for 16 hours after which medium was changed back to growth medium. After 24 and 48 hours in growth medium, supernatant containing lentivirus was collected and spun at 9,000 rpm for 10 minutes to remove cell debris. 50% polyethylene glycol (w/w) was applied to the supernatant in the ratio of 1 to 8 respectively and incubated for 16-20 hours at 4°C to precipitate lentivirus. Supernatant was then again centrifuged at 9,000 rpm for 15 minutes at 4°C and pellet was suspended in cold D-PBS, filtered through 0.45 µm PVDF filter, and centrifuged at 21,000xg for 1 hour at 4°C. Pellets containing lentivirus were resuspended in 20µL of D-PBS (for 6 dishes), vortexed, aliquoted, and stored at -80°C until further use. Lentiviral titer (transducing units/µL) was calculated by serially diluting (10 times) lentivirus on plated 50,000 U251 cells per well with 8 µg/ml of hexadimethrine bromide or polybrene (H9268-5G, Sigma) and quantifying the total number of infected cells with that lentiviral dilution per well that lie within the dynamic range of 1-10% infectivity.

### Generation of stable monoclonal U251 cell lines

50,000 U251 cells were plated in growth medium in each well of a 24-well plate (353047, Falcon). Lentivirus expressing either WT αSyn-EGFP or A53T αSyn-EGFP or A53T αSyn-mTurquoise2-IRES-A53T αSyn-mNeonGreen was applied to each well with a multiplicity of infection (MOI) equaling to 30. Lentivirus treated cells were expanded to 10 cm dish until confluency was reached. BD FACSMelody (BD Biosciences) cell sorter was used to select 10% brightest EGFP positive single cells (excitation 488 nm laser and emission 527/32 nm) expressing WT αSyn-EGFP or A53T αSyn-EGFP. BD FACSymphony S6 (BD Biosciences) cell sorter was used to select 5% brightest A53T αSyn-mNeonGreen positive single cells (excitation 488 nm laser and emission 515/20 nm) co-expressing A53T αSyn-mTurquoise2 (excitation 405 nm laser and emission 450/50 nm). Unstained U251 cells and cells expressing single fluorophore were used as compensation controls to correct fluorescence spillover. 200 single cells were plated in 100 µL fresh growth medium per well of a 96-well plate (P96-1.5P, CellVis). 20 colonies were selected based on growth and expanded in a 6-well plate (3516, Corning Costar). To identify a U251 monoclonal cell line out of 20 colonies that allows efficient aggregation of intracellular αSyn by exogenous recombinant αSyn pre-formed fibrils (rPFF), 10,000 cells from each cell clone was plated in triplicates in each well of 96-well plate and 3 µM of sonicated rPFF was applied to the cells 1 day after plating. 4 days post-application (DPA) of rPFF, which is 5 days *in vitro* (DIV), detergent soluble fraction was extracted and removed from the cells with cold 4% sucrose and 1% nonidet-40 (NP-40 or IGEPAL) in 1X D-PBS for 2 minutes. Immediately after, cells were fixed with cold 4% paraformaldehyde (PFA) (R04586-76, Sigma) and 1% NP-40 in 1X D-PBS for 15 minutes. The monoclonal cell line with the highest number of detergent insoluble puncta was selected, when excited with 482/25 nm LED and detected with 524/24 nm filter set in Evos M7000 (Thermo Scientific), expanded further, and frozen in liquid nitrogen with 10% DMSO in growth medium.

### Rapid *in situ* detection of fluorescent intracellular αSyn aggregates

Fluorescent αSyn aggregates formed intracellularly by seeding with rPFF were rapidly detected either in U251 cells expressing WT αSyn-EGFP or A53T αSyn-EGFP, or U251 biosensor cells expressing A53T αSyn-mTurquoise2-IRES-A53T αSyn-mNeonGreen. Culture supernatant was discarded and cells were treated with cold 4% sucrose and 1% NP-40 in 1X D-PBS for 2 minutes. The sucrose-NP-40 solution was discarded and cells were immediately fixed with cold 4% PFA and 1% NP-40 in 1X D-PBS for 15 minutes. While the cells were being fixed, fluorescent αSyn aggregates resembling ring-like structures or puncta, were easily detected in Evos M7000 when excited with 482/25 nm LED and detected with 524/24 nm filter set. After fixing, cells were rinsed once with 100 µL of D-PBS and stored in 200µL of D-PBS containing 0.03% sodium azide (storage buffer) at 4°C. Intracellular fluorescent αSyn aggregates were more clearly visible in storage buffer.

### Intracellular αSyn aggregation induced by exogenous proteopathic seeds

For “naked” seeding with recombinant αSyn pre-formed fibrils (rPFF), monoclonal U251 cell line expressing WT αSyn-EGFP or A53T αSyn-EGFP, 10,000 cells were plated in each well of 96-well plate in growth medium. Next day, 100% medium was changed to 90 µL of fresh medium. Required volume of rPFF, at least 3 µL in final volume of 15 µL D-PBS, was prepared in 1.5 ml polystyrene tubes was placed in a water bath pre-set to 4°C in a cup-horn sonicator (Q700, 431C2, QSonica). 10 minutes of sonication was set and carried out at an amplitude of 70 with 30 seconds of sonication and 30 seconds of rest with 250-350 W power output per cycle. 10 µL of sonicated rPFF per well were applied to the cells within 2 hours of sonication so that the final concentration per well was 3 µM (4,320 ng), which was equivalent to 432 pg per cell. D-PBS vehicle or αSyn monomer equivalent in D-PBS was used as a negative control.

For monoclonal biosensor U251 cell line expressing A53T αSyn-mTurquoise2 and A53T αSyn-mNeonGreen, 30,000 cells were plated in each well of 96-well plate in growth medium. Next day, 100% medium was changed to 120 µL of fresh medium. Required volume of skin or brain-amplified fibrils, at least 3 µL in final volume of 15 µL D-PBS, was prepared in 1.5 ml polystyrene tubes and sonicated as described above. Sonicated patient-amplified αSyn fibrils were diluted further in cold D-PBS as desired. 6% (v/v) lipofectamine 2000 (11668027, Invitrogen) in Opti-MEM (31985070, Gibco) was incubated for 5 minutes at room temperature. Equal volume of sonicated patient-amplified αSyn fibrils was applied to 6% lipofectamine drop-wise and gently mixed by triturating 4-5 times. Lipofectamine-αSyn fibril mix was incubated for at least 30 minutes at room temperature. 30 µL of lipofectamine-αSyn fibril mix was applied to each well of cells containing 120 µL of growth medium. Final concentration of lipofectamine 2000 in each well of 96-well plate was 0.6% in 150 µL final medium volume. Final concentration of patient-amplified αSyn fibrils per well of 96-well plate was 100 nM (216 ng), which was equivalent to 7.2 pg of patient fibrils per cell. Concentrations of patient-amplified αSyn fibrils and lipofectamine were kept constant while using a 6-well plate, wherein, 180 µL of Lipofectamine-αSyn fibril mix was applied to 200,000 cells in a final medium volume of 1 ml per well resulting in 100 nM (1,440 ng) of patient-derived fibrils per well, which was also equivalent to 7.2 pg of patient fibrils per cell. Equimolar concentrations of lipofectamine-αSyn monomer and lipofectamine-rPFF mix were used as controls. Experiments were carried out until desired time-point with medium replenishments and cells were processed accordingly for further analyses by RT-QuIC assay, immunofluorescence, immunoblotting, and FRET-Flow cytometry.

### FRET-Flow cytometry

100 nM of skin- or brain-amplified αSyn fibrils with lipofectamine were applied on 30,000 monoclonal biosensor U251 cells in a well of a 96-well plate as described above. After 4 days post application (DPA) of αSyn fibrils, supernatant was discarded, and cells were rinsed once with 50 µL D-PBS per well. Cells were incubated at 37°C with 50 µL of 0.05% trypsin for 3 minutes that was deactivated with 130 µL of growth medium. Dissociated cells were transferred into 96-well U-bottom plate (7007, Corning) and spun at 1,000xg for 3 minutes. Medium was carefully discarded, and cells were suspended and fixed in 50 µL of cold 2% PFA in D-PBS for 10 minutes at room temperature. After fixing, plate was spun at 1,000xg for 3 minutes and cell pellet was resuspended in 105 µL of cold FACS buffer containing 2% FBS and 1mM EDTA in D-PBS. 70 µL of biosensor cell suspension was aspirated by Attune NXT flow cytometer (ThermoFisher) and FlowJo v10.10.0 software was used to quantify percent of cells positive for FRET (FRET percentage) and efficiency of αSyn oligomerization in cells (FRET efficiency). Median fluorescence intensities of mTurquoise2 (donor) and FRET were measured by exciting U251 cells at 405 nm and detecting emission at 440/50 nm and 530/30 nm respectively. Median fluorescence intensity (MFI) of mNeonGreen (acceptor) was measured by exciting cells at 488 nm and detecting emission at 530/30 nm. MFI ratio of donor to acceptor in monoclonal biosensor cell line was found to be 1 to 8.9 ± 0.3 SEM (N = 12). To quantify FRET, gating strategy was adapted from previously described study [60]. Bleedthrough of mNeonGreen into FRET channel was negligible in these cells. A bivariate plot of mTurquoise2 and FRET was created. Lipofectamine-αSyn monomer treated cells were used to adjust gate to exclude single cell population that do not contain αSyn aggregates and were thus FRET negative. This gate on the bivariate plot allowed us to quantify FRET positive cells with sensitized acceptor emission at 530/30 nm by donor excitation at 405 nm occurring due to αSyn oligomerization and formation of intracellular αSyn aggregates. Integrated MFI was calculated by the product of FRET percentage and FRET MFI. Finally, to measure the efficiency of αSyn oligomerization events, FRET efficiency was calculated and reported by the following formula:

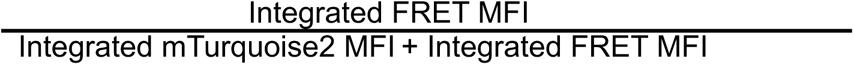

Each data point resulted from 20,000 to 25,000 single cells analyzed out of 100,000 to 110,000 total cells per well.

### Immunofluorescence

100 nM of skin- or brain-amplified αSyn fibrils with lipofectamine were applied on 30,000 monoclonal biosensor U251 cells in a well of a 96-well plate as described above. After 4 DPA of αSyn fibrils, supernatant was discarded, and cells were rinsed once with 50 µL D-PBS. Cells were then fixed in 100 µL of cold 4% PFA for 15 minutes at room temperature. Cells were rinsed twice with 100 µL of cold D-PBS, and stored in 200 µL of D-PBS with 0.03% sodium azide at 4°C. Immunofluorescence was performed as previously described [61,62]. Storage solution was discarded, and cells were blocked in blocking buffer containing 5% normal donkey serum (0030-01, SouthernBiotech) in 0.3% NP-40 in 1X D-PBS (rinse buffer) for 20 minutes at room temperature in dark. Primary antibody was diluted in fresh blocking buffer and incubated with cells at room temperature for 1 hour in dark. Cells were rinsed 3 times in 100 µL of rinse buffer followed by incubating with secondary antibody diluted in fresh blocking buffer at room temperature for 1 hour in dark. To counterstain nuclei (if required), 50 µL of 4 µg/ml of DAPI (D9542, Sigma) in D-PBS was applied to the cells and incubated for 3 minutes after discarding secondary antibody solution. Cells were then rinsed 3 times in 100 µL of rinse buffer and once with 1X D-PBS followed by storing in 200µL of fresh D-PBS. Plate was then sealed with Nunc clear sealing film and was stored with lid closed at 4°C. Primary antibodies used were as following: rabbit anti-p-αSyn (1:500 dilution, EP1536Y, Abcam), mouse anti-TUJ1 (1:1000, MAB1195, R&D Systems), mouse anti-NeuN (1:500, MAB377, Abcam), and mouse anti-NEUROD (1:250, sc-46684, Santa Cruz). Secondary antibody used were as following: donkey anti-rabbit IgG Alexa fluor 647 (1:500, A-31573, Invitrogen), donkey anti-mouse IgG Alexa fluor 555 (1:500, A-31570, Invitrogen), and donkey anti-mouse IgG Alexa fluor 647 (1:500, A-31571, Invitrogen).

### Fluorescence microscopy and imaging

Fluorescent images were acquired either by Nikon A1R-HD Confocal T-2 Eclipse Inverted Microscope with Nis Elements software using Gasp PMT Detectors (Fig. 2, 3) or Leica Stellaris 5 Confocal DMI8 Inverted Microscope with LAS-X Software using HyD Detectors (Fig. 5) or BioTek Cytation C10 spinning-disk confocal imaging reader with Gen5 software (Agilent) (Fig. 1, 6). On Nikon A1R-HD Confocal, FRET was excited at 409 nm and its emission was detected by 538/42 filter set. mTurquoise2 was excited at 409 nm and its emission was detected by 450/25 filter set. DAPI was excited at 409 nm and its emission was detected by 450/50 filter set. EGFP and mNeonGreen were excited at 489 nm and their emission was detected by 525/50 filter set. Alexa Fluor 647 bound to p-αSyn antibody was excited at 638 nm and its emission was detected by 700/75 nm filter set. On Leica Stellaris 5 Confocal microscope, FRET was excited at 405 nm and its emission was detected by 532/37 filter set. mTurquoise2 was excited at 405 nm and its emission was detected by 455/41 filter set. Alexa Fluor 647 bound to p-αSyn antibody was excited at 653 nm and its emission was detected by 709/100 nm filter set. mNeonGreen was excited at 488 nm and its emission was detected by 525/50 filter set. To counterstain nuclei, SYTOX orange was excited at 553 nm and its emission was detected by 600/84 filter set. On BioTek Cytation C10 spinning-disk confocal reader, DAPI was excited at 377 nm and its emission was detected by 447/60 nm filter set. FRET was excited using 400/40 nm LED and its emission was detected by 550/49 nm filter set. EGFP was excited at 472 nm and its emission was detected by 525/39 nm filter set. SYTOX orange was excited at 556 nm and its emission was detected by 600/37 filter set. Alexa Fluor 647 bound to p-αSyn antibody was excited at 635 nm and its emission was detected by 685/40 nm. When live cells were analyzed on Evos M7000, FRET was excited by 445/45 nm LED and its emission was detected by 542/27 nm filter set. mNeonGreen was excited by 482/25 nm LED and its emission was detected by 524/24 nm filter set. dsRed was excited by 531/40 nm LED and its emission was detected by 593/40 filter set.

**Figure 1:**
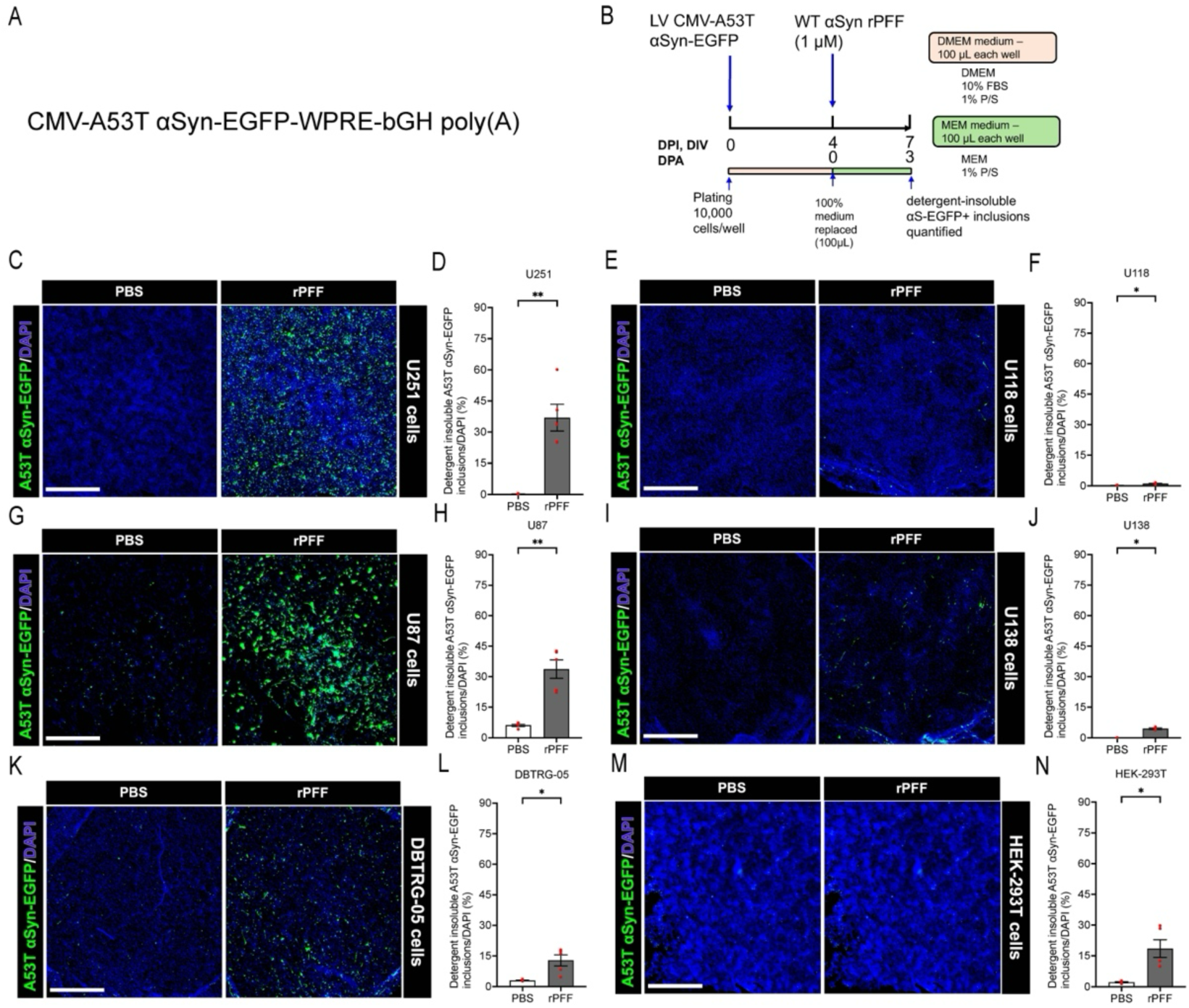
Human U251 glioblastoma cell line allows robust aggregation of intracellular A53T αSyn-EGFP by naked seeding with exogenous WT αSyn rPFF. (A) The lentiviral vector for expression of A53T αSyn-EGFP. (B) Experimental design showing the timelines of lentivirus and αSyn rPFF application along with medium changes. (C-N) Detergent-insoluble A53T αSyn-EGFP inclusions detected in the indicated cell lines 3 after application of αSyn rPFF. Cells were analyzed after *in situ* detergent extraction of soluble proteins, followed by immediate fixation with 4% PFA. D-PBS was used as a vehicle for all experiments. A53T αSyn-EGFP aggregates are shown in green (puncta) whereas nuclei are shown in blue. Quantification of detergent-insoluble A53T αSyn-EGFP aggregates (D, F, H, J, L, N). Data are presented as mean ± SEM, where each dot corresponds to a biological replicate in which (D) 21,200, (F) 14,812, (H) 6,374, (J) 10,330, (L) 12,524 cells on average were analyzed; N=4-5 per group. Scale bars: 1 mm. Statistical analysis was performed by unpaired two-tailed Student’s t-test with Welch’s correction or unpaired two-tailed Mann-Whitney test (D, J), and shown as *p<0.05, **p<0.01, ***p<0.001, ****p<0.0001. EGFP: enhanced green fluorescent protein, LV: lentivirus, WT αSyn rPFF: wild-type alpha-synuclein recombinant pre-formed fibrils, DIV: days *in vitro*, DPA: days post-application of rPFF, DPI: days post-infection with lentivirus.

**Figure 2:**
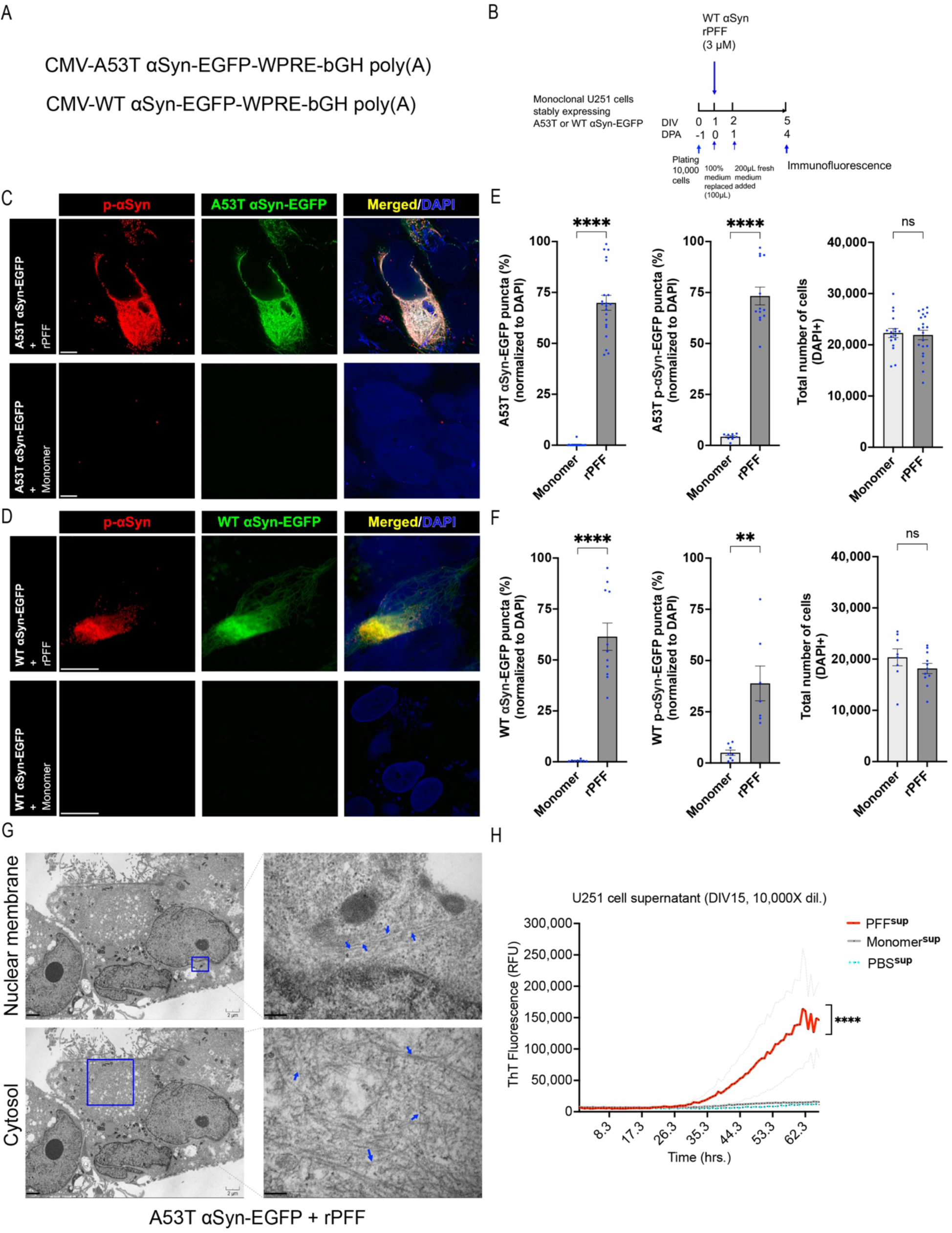
Naked seeding with rPFF results in pathological αSyn inclusions in U251 cells that release proteopathic αSyn seeds. (**A**) The lentiviral vectors for expression of A53T αSyn-EGFP or WT αSyn-EGFP in U251 cells. (**B**) Experimental design showing the timelines of lentivirus and αSyn rPFF or αSyn monomer application. Monoclonal cell line selection was performed using flow cytometry to select a cell line that has the highest naked seeding efficiency induced by αSyn rPFF. (**C, D**) Representative confocal images showing pathological detergent-insoluble A53T or WT αSyn aggregates (green) that are phosphorylated (red). Cells were analyzed at DIV 5 after *in situ* extraction of soluble proteins. Scale bars: 5 µm. (**E, F**) Quantification of cells with detergent-insoluble WT or A53T αSyn aggregates. Data are presented as mean ± SEM, and each dot corresponds to a biological replicate where 15,000 to 25,000 cells were analyzed; N=3-4 per group from 4-5 independent experiments (E) and N=3 per group from 2-3 independent experiments (F). (**G**) Transmission electron microscopy (TEM) images of cellular sections showing fibrillar assemblies around nuclear membrane and in cytosol (blue arrows) in cells expressing A53T αSyn-EGFP and treated with rPFF. Scale bars: 2 µm for the lower magnification images, 0.2 µm for the higher magnification nuclear membrane image, and 0.5 µm for the higher magnification cytosol image. (**H**) Pathologic αSyn seeds detected in the supernatant of cells treated with rPFF, but not in αSyn monomer treated supernatant, by real-time quaking-induced conversion (RT-QuIC) measured as a function of ThT fluorescence. Data are presented as mean ± SEM (grey line), and each dot represents data from N=2-5 replicates per group. Statistical analysis was performed by unpaired two-tailed Mann Whitney test or unpaired two-tailed student’s t-test, and Welch’s correction was applied when variances were found to be significant. A p-value <0.05 was considered significant. Significant differences are indicated by *p<0.05, **p<0.01, ***p<0.001, ****p<0.0001.

### Purification of intracellular αSyn fibrils from U251 biosensor cells

300 nM of skin- or brain-amplified αSyn fibrils with lipofectamine were applied on 200,000 monoclonal biosensor U251 cells in a 6-well plate as described above. After 5 DPA of αSyn fibrils, supernatant was discarded and 250 µL of cold extraction buffer containing 50 mM Tris pH 7.4 at 25°C (T-2663, Sigma), 800 mM NaCl, 1% NP-40, 10% sucrose (w/v), 1X EDTA-free protease and phosphatase inhibitor (A32961, Thermo), 1 mM EDTA, and 1 mM DTT was applied on the cells and incubated for 2 minutes at room temperature to extract detergent-soluble proteins. The applied extraction buffer was saved as detergent soluble fraction. Then, NP-40 in extraction buffer was replaced by 2% sarkosyl (61747, Sigma) and was applied to the cells. Cells were scrapped in 300 µL of sarkosyl buffer and plate was nutated at room temperature for 1 hour. Cell homogenate was transferred to 1.5 ml tube, nutated at 37°C for 15 minutes, and then sonicated at amplitude 30 for 30 seconds at 4°C. Homogenate was spun at 10,000xg for 10 minutes at 4°C to pellet out cell debris. Supernatant was collected and spun at 186,000xg for 30 minutes at 4°C. Supernatant was collected and mixed with detergent soluble fraction that was collected earlier. Pellet was rinsed with 800 µL of cold D-PBS followed by spinning at 186,000xg for 30 minutes. Pellet was resuspended in 15 µL of cold D-PBS. Protein concentration was measured using Nanodrop A_205_ as previously described. For RT-QuIC analysis and amplification of U251 cell-derived αSyn fibrils, the detergent-insoluble fraction was sonicated at amplitude 70 for 10 minutes with 30 seconds sonication and 30 seconds rest at 4°C as previously described. Sonicated fraction was then used for RT-QuIC amplification or stored at -80°C for later use. For immunoblotting, 4X LDS sample buffer (B0007, Novex) with 200 mM DTT was applied to the detergent insoluble fraction (1.2X LDS-DTT sample buffer final concentration), heated to 95°C for 5 minutes, and stored at -20°C for further analysis.

### Immunoblotting

Immunoblotting was performed as previously described [58,61]. 4X LDS sample buffer with 200 mM DTT was applied to detergent soluble or insoluble fraction or clarified cell lysate (1.2X LDS-DTT sample buffer final concentration). Samples were heated to 95°C for 5 minutes. Samples equivalent to 5 µg total protein amount were loaded on 4-12% bis-Tris protein gel (NP0323BOX, Fisher) and run in 1X Tris-MES-SDS running buffer (pH 7.3) containing 50 mM Tris base, 50 mM MES, 0.1% SDS, and 1 mM EDTA at 100 V for 1.5 hours. Protein bands were transferred to 0.45 µm pore sized PVDF membrane (IPVH00010, Immobilon-P, Millipore) in cold 1X Tris-glycine transfer buffer containing 25 mM Tris, 192.4 mM glycine, and 20% methanol at 20 V (2.5 A limit) for 1 hour by a semi-dry method using Trans-Blot Turbo transfer system (Bio-Rad). After transfer, PVDF membrane was rinsed once in 0.1% tween-20 in D-PBS (wash buffer) and blocked in 5% BSA in 0.1% tween-20/D-PBS (block buffer) for 20 minutes. Primary antibody was diluted in block buffer and applied to PVDF membrane that was incubated at 4°C in dark for 16 hours. PVDF membrane was then rinsed 3 times in wash buffer. Secondary antibody coupled to horseradish peroxidase was diluted in block buffer and then applied to PVDF membrane that was incubated at room temperature in dark for 1 hour. PVDF membrane was again rinsed 3 times in wash buffer. ECL (RPN2232, Cytiva) was used for chemiluminescent detection of protein bands using ChemiDoc imaging system (Bio-Rad). Primary antibodies used were as following: rabbit anti-αSyn (1:1000 dilution, MJFR1, Abcam), rabbit anti-p-αSyn (1:1000, EP1536Y, Abcam), and mouse anti-GAPDH (1:10,000, 60004-1-Ig, Proteintech). Secondary antibodies used were as following: donkey anti-rabbit HRP (1:10,000, SA1200, Thermo Scientific), and donkey anti-mouse HRP (1:10,000, SA1100, Thermo Scientific).

### Quantification and classification of p-αSyn aggregate morphology

Pathological αSyn (p-αSyn positive) aggregates in biosensor U251 cells and induced biosensor neurons were quantified and their morphology was assessed post fixation in 4% PFA. 96-well plate was loaded into BioTek Cytation C10 spinning-disk confocal imaging reader (Agilent), and confocal images were captured by Hamamatsu scientific CMOS Orca camera and automatically deconvoluted by Gen5 software (Agilent). Biosensor U251 cell nuclei were counterstained with 0.5 µM of SYTOX Orange (S11368, Invitrogen) in D-PBS after imaging p-αSyn aggregates, whereas mouse anti-TUJ1 antibody was used to counterstain induced biosensor neurons simultaneously with rabbit anti-p-αSyn antibody. Nuclei and cell bodies (soma with nucleus) were quantified automatically using Gen5 software with optimized settings. Intracellular p-αSyn aggregates in biosensor U251 cells were classified into 4 classes of phenotypic morphologies based on mean fluorescence intensity (MFI) and size of the p-αSyn aggregates. They were named diffuse speckles, diffuse inclusions, compact speckles, and compact inclusions. MFI of 8,000 to 14,999 was termed as diffuse, whereas MFI ≥15,000 was termed as compact. When a p-αSyn aggregate cover a smaller area of 0.5-4.5 µm in a cell, it was termed as a speckle, whereas, when it covers a larger area of 4.6-35 µm, it was termed as an inclusion. p-αSyn aggregates in biosensor cells that meet the above criteria were quantified automatically using Gen5 software. In U251 biosensor cell-derived induced neurons, only those p-αSyn aggregates that overlap specifically with TUJ1 positive neuronal cell bodies and meet the above criteria were quantified automatically. Length of p-αSyn aggregates in neuronal cell bodies were calculated automatically using Gen5 software by drawing a straight line across the farthest points in those aggregates. Images were acquired from 4 different locations in a well of 96-well plate. Precisely, same location in all wells were used for image acquisition. Total area of all locations in a well amounted to 3.86 mm^2^ for analyzing biosensor cells and 8.06 mm^2^ for induced biosensor neurons. Each data point was represented as mean ± SEM of all four images from each well. Quantified p-αSyn aggregates were normalized to the area and number of nuclei or cell bodies from each image in each well. Proportion of each morphology for each disease group in biosensor cells was quantified after normalization and plotted against each morphological phenotype of speckles and inclusions. From biosensor cells, each data point resulted from 5,000 to 10,000 cells with 1,500 to 14,000 p-αSyn speckles and inclusions analyzed. From induced biosensor neurons, each data point resulted from 180 to 700 neuronal cell bodies analyzed.

### Generation of U251 induced biosensor neurons

Generation of induced neurons from U251 biosensor cells was adapted from a prior study [63]. Monoclonal biosensor U251 cells were transduced with lentivirus to deliver NEUROD1 and dsRed or dsRed alone (control) with a MOI equaling to 15. Lentivirus that delivered dsRed only was used a control for reprogramming. Lentivirus was applied to cell suspension containing growth medium with 8 µg/ml polybrene for 15 minutes in 37°C water bath with intermittent mixing. 2,700 cells in 200 µL of growth medium were plated in each well of a 96-well plate, coated with Matrigel (354277, Corning), which was diluted 500 times in DMEM. After 3 days post infection (DPI), which was considered days *in vitro* (DIV) 0, 100% of growth medium was replaced with neuronal differentiation medium containing DMEM/F12/Neurobasal (2:2:1) (SH30243 Cytiva, 51445C Sigma, 12348017 Gibco), 0.8% N2 (17502048, Gibco), 0.4% B-27 plus (A3653401, Gibco), with small molecules 1 µM dorsomorphin (P5499, Sigma), 10 µM forskolin (S2449, Selleck Chemicals), with growth factors 20 ng/ml BDNF (450-02, Gibco), 20 ng/ml GDNF (450-10, Gibco), 20 ng/ml NT3 (450-03, Gibco), and 1% P/S. 50% neuronal medium with freshly added small molecules and growth factors was changed every other day until 21 DPI after which medium replacement was done twice a week. Lentivirus transduction was reported by dsRed positive cells. NEUROD1 expressing cells develop long neuritic extensions emerging from the cell body causing dsRed levels to diffuse resulting in lower dsRed signal intensity. Control group of cells do not express NEUROD1 and thus, do not develop long neurites from the cell body thereby accumulating dsRed signal with relatively higher intensity in the cell body. 100 nM of skin or brain-amplified αSyn fibrils with lipofectamine was applied on DPI 34 (DIV 31), as described above, and replaced with 200 µL of fresh neuronal medium after 16 hours. αSyn monomer equivalent with lipofectamine was used as a control. After 14 days post application (DPA) of patient fibrils (DPI 48, DIV 45), human induced biosensor neurons (hiBNs) were processed for desired analyses.

### Electrophysiology of induced biosensor neurons

14,000 monoclonal biosensor U251 cells transduced with lentivirus delivering NEUROD1 and dsRed were plated on Matrigel coated coverslips (6330029, Carolina Science and Math). At DPI 48 (DIV 45), these coverslips were transferred to a chamber on a BX51WI Olympus microscope and superfused with recording solution containing 125 mM NaCl, 2.5 mM KCl, 1.25 mM Na_2_HPO_4_, 1 mM MgCl_2_, 2 mM CaCl_2_, 25 mM D-glucose and 25 mM NaHCO_3_ bubbled with 5% CO_2_/95% O_2_. dsRed reporter was used to track induced biosensor neurons and were also visualized using epifluorescence illumination with Texas Red or FITC filter sets. Fire-polished borosilicate glass electrodes (BF150-86-10, Sutter Instrument) with resistance of 4–6 MΩ when filled with intracellular solution were mounted on the headstage (CV-7B) of a Multiclamp 700A amplifier (Molecular Devices). For whole-cell recordings, patch pipettes were filled with 135 mM K-gluconate, 2 mM MgCl_2_, 10 mM HEPES, 3 mM KCl, 0.5 mM Na-GTP, 2 mM Mg-ATP, 10 mM phosphocreatine, and 0.1 mM EGTA, pH 7.3, and 310 mOsm. Recordings were made at room temperature and bridge balance was automatically adjusted in current-clamp recordings to compensate series resistance (<25 mΩ). Currents were sampled at 10 kHz and filtered at 2 kHz (Digidata 1440A; Molecular Devices) using PClamp 10 software (Molecular Devices). Action potentials were elicited in current clamp by increasing current steps measured during 500 ms. Sodium currents induced by increasing voltage steps (300 ms) were measured in cells held at -80 mV. Data were analyzed with Clampfit (Molecular Devices).

### Statistical analyses

All statistical analyses were performed using GraphPad Prism v10.4.1 (GraphPad). All data were examined for normality by Shapiro Wilk test. For non-normally distributed data, unpaired two-tailed Mann-Whitney’s test was used for two groups, and Kruskal-Wallis with Dunn’s multiple comparison test was used for three or more groups. All data were also examined for differences in variance. For significantly different variance among groups, unpaired two-tailed Student’s t-test with Welch’s correction was used for two groups, and Brown-Forsythe and Welch ANOVA with Dunnett’s T3 multiple comparison test was used for three or more groups. For normally distributed data with no significant differences in variance, unpaired two-tailed Student’s t-test was used for two groups, and one-way or two-way ANOVA with Tukey’s multiple comparison test was used for three or more groups. Null hypothesis was rejected if P value was < 0.05, which was therefore considered statistically significant. Statistical powers of comparisons were analyzed by G*Power 3.1 [64] and values were reported on the plots. Following settings were used: t-tests, two-tailed, difference between two independent means (two groups), post hoc power analysis, and an α error probability of 0.05. The effect size d was determined by means and standard deviation of the two groups that were being analyzed, and the sample size of the two groups were provided. Using the settings with determined and provided values, the power (1-ß error probability) of the statistical assessment was calculated. Statistical power of the respective statistical analysis was accepted to be reasonable if it was greater than 0.8.

## Results

### Human U251 glioblastoma cells allow robust aggregation of intracellular αSyn upon “naked” seeding with rPFF

Intracellular milieu modulates αSyn aggregation and is one of the major factors that is responsible for generation of different strains of αSyn in Lewy body diseases and multiple system atrophy [46,47]. Different cell models have been developed to study intracellular αSyn protein aggregation trigged by exogenous proteopathic αSyn “seeds”, a process also known as “seeding” [65–68]. However, formation of intracellular αSyn aggregates that are detergent-insoluble and occur homogenously in most cells upon exogenous αSyn seeding remain unachieved. Furthermore, most of the cell models are not as robust as they are difficult to grow and maintain in culture and may require labor- and time-intensive differentiation protocols.

To identify a cell line that robustly aggregates intracellular αSyn by naked seeding with αS proteopathic seeds and is easily maintained in culture, we tested different glioblastoma cell lines because they are derived from human brain and have negligible levels of endogenous αS [69] which is known to interfere with αSyn seeding and aggregation in engineered cells [70]. Human αSyn (encoded by SNCA gene) carrying the disease-causing A53T mutation was tagged with EGFP (A53T αSyn-EGFP) (Fig. 1A) and was constitutively expressed in human glioblastoma cell lines U251, U118, U87, U138, DBTRG-05, and a human embryonic kidney (HEK) 293T cell line using lentivirus-mediated transduction. Freshly sonicated recombinant αSyn pre-formed fibrils (rPFF), commonly used as proteopathic αSyn seeds [27,28] was applied to the medium of cells at 1 mM (Fig. 1B). Treatment of cells with PBS was used as a control. 3 days post application (DPA) of rPFF seeds, soluble proteins were removed from the cells by *in situ* extraction with detergent resulting in bright detergent-insoluble intracellular A53T αSyn-EGFP positive inclusions visualized by fluorescence microscopy (Fig. S1). A53T αSyn-EGFP inclusions were normalized to the total number of cells (Fig. S2) and percentage of detergent-insoluble αSyn inclusions were reported. Among all cells tested, U251 cell line showed highest percentage of detergent-insoluble αSyn inclusions induced by exogenous rPFF seeds followed by U87, HEK-293T, DBTRG-05, U138, and U118 cell line (Fig. 1C-N). Furthermore, U251 cell line reported highest total number of cells survived after 3 DPA of rPFF, followed by HEK-293T, U118, DBTRG-05, U138, and U87 (Fig. S2). rPFF showed similar cell survival as compared to the PBS control (Fig. S2). Overall, our results suggest that human U251 glioblastoma immortalized cell line provides an ideal cellular milieu for intracellular αSyn aggregation upon seeding by rPFF under quasi-physiological conditions, and it allows rapid *in situ* detection of detergent-insoluble intracellular αSyn inclusions.

### Naked seeding with rPFF result in pathological phospho-αSyn inclusions in U251 cells that release proteopathic αSyn seeds

Intracellular αSyn aggregation can be induced by rPFF seeds prepared with different sonication parameters and instruments and in different culture media, which may contribute to the variability in the reported seeding efficiency of rPFF in cell models [27,46,66,68,71–75]. Therefore, cell culture conditions were optimized for enhancing the number of detergent-insoluble αSyn inclusions in U251 cells upon seeding with rPFF (Fig. S3). Serum concentration and instrument-specific sonication parameters were observed to affect rPFF seeding efficiency in U251 cells (Fig. S3C, H). A dose-dependent increase of intracellular αSyn aggregates detected as fluorescent puncta was observed with increasing concentration of exogenous rPFF in the medium (Fig. S3J). To examine if the detergent-insoluble inclusions induced by rPFF are biochemically and pathologically similar to those present in diseased brains, immunostaining was performed to detect phospho-serine129 in αSyn (p-αSyn), a well-known maker for pathological αSyn aggregates in Lewy bodies and Lewy neurites in synucleinopathies [6,7]. Monoclonal U251 cell lines were generated that constitutively express either A53T αSyn-EGFP or WT αSyn-EGFP (Fig. 2A). Immunofluorescence of p-αSyn was performed 4 days after rPFF application on U251 cells, followed by *in situ* removal of soluble proteins with detergent extraction (Fig. 2B). Cells treated with αSyn monomer were used as control. p-αSyn was strongly detected in the detergent-insoluble A53T or WT αSyn-EGFP positive inclusions (Fig. 2C, D) that were quantified (Fig. 2E, F). Detergent-insoluble and phosphorylated αSyn inclusions were around 2.3-fold higher in A53T αSyn-EGFP expressing U251 cells in comparison to the WT αSyn-EGFP expressing cells. Total number of cells used for analysis in the control and rPFF groups were not significantly different (Fig. 2E, F). Our results suggest that seeding by exogenous rPFF can trigger aggregation of intracellular A53T and WT αSyn-EGFP to form pathological p-αSyn inclusions.

Next, to examine the subcellular localization of detergent-insoluble αSyn species, TEM of ultrathin sections of U251 cells with A53T αSyn inclusions trigged by rPFF seeding was performed. TEM images revealed fibrillar assemblies in the cytosol and around the nuclear membrane (Fig. 2G, blue box), consistent with the subcellular localization of p-αSyn positive inclusions (Fig. 2C). Furthermore, the width of these fibrillar assemblies in cells ranged from 9.9-10.7 nm in width (Fig. 2G, blue arrows). These assemblies were not detected in the αSyn monomer treated control group (Fig. S4A).

Finally, we investigated if U251 cells containing intracellular αSyn inclusions can release pathological αSyn seeds. rPFF seeds were applied to monoclonal U251 cells expressing A53T αSyn-EGFP at day 1. After several medium changes, cell culture supernatants were collected and immunofluorescence of cells was performed at day 15 (Fig. S4B). αSyn monomer and PBS treated cell supernatants were used as controls. Detergent-insoluble A53T αSyn-EGFP inclusions were detected only in rPFF-treated U251 cells and quantified (Fig. S4C-D). To examine whether these U251 cells release pathological αSyn seeds capable of inducing seeding activity *in vitro*, an ultrasensitive seed amplification assay known as RT-QuIC was performed with the cell supernatants. RT-QuIC assay of 10,000x diluted supernatants collected from U251 cells expressing A53T αSyn-EGFP treated with rPFF readily revealed strong αSyn seeding activity with peak ThT fluorescence at around 60 hours, but not in the cells treated with αSyn monomer or PBS control supernatants (Fig. 2H). These data suggest that αSyn-inclusion-containing U251 cells appear to release pathological αS seeds. Taken together, our results indicate that rPFF seeds trigger the formation of pathological p-αSyn-positive inclusions in U251 cells that are detergent-insoluble and are localized in the cytosol and around the nuclear membrane. The U251 cells containing these αSyn inclusions release proteopathic αSyn seeds.

### Generation of FRET-based αSyn U251 biosensor cell line

To study intracellular αSyn aggregation in live cells in response to exogenous proteopathic seeds, FRET-based αSyn biosensors were developed with U251 cells. Previously, HEK-293T FRET-based biosensor cells with donor/acceptor fluorophore pair or with single fluorophores have been widely used to detect αSyn and tau aggregation [71,72]. Monocistronic plasmids expressing either donor or acceptor fluorophore [71,76] might result in different proportions of donor and acceptor protein expression in each cell, thereby attributing to variability in FRET readout. Here we designed a bicistronic vector with both donor and acceptor transgenes. CMV promoter controlled the expression of mTurquoise2 donor fused to the C-terminus of A53T αSyn (αSyn-mTQ2/αSyn-CFP), in tandem with the IRES controlled expression of mNeonGreen acceptor fused to the C-terminus of A53T αSyn (αSyn-mNG/αSyn-YFP) (Fig. 3A). Monocistronic plasmids expressing either αSyn-mTQ2 or αSyn-mNG were also constructed and used as negative controls (Fig. 3A). A total of 192 single U251 cells constitutively expressing αS-mTQ2 and αS-mNG were isolated by flow cytometry and a single cell clone was selected based on its efficiency to aggregate A53T αSyn into detergent-insoluble puncta upon naked seeding by rPFF seeds. The selected FRET monoclonal biosensor U251 cell clones were then plated along with control cell lines expressing either of the two fluorophores and rPFF seeds were applied (Fig. 3B). Monomeric αSyn was used as a treatment control. 5 days after application of rPFFs, FRET-Flow cytometry was performed (Fig. 3B). Strategically, a subset of single-cells expressing αSyn-mNG were always selected from single-cell population that were also found to be expressing αSyn-mTQ2 (Fig. S5A). Control cell line groups with monomeric αSyn treatment expressed either αSyn-mTQ2 or αSyn-mNG, whereas biosensor cells with αSyn monomer or rPFF treatment express both αSyn-mTQ2 and αSyn-mNG (Fig. S5B). Control and biosensor cells treated with αSyn monomer were excited to detect bleed-through of either αSyn-mNG or αSyn-mTQ2 into FRET channel using which FRET gates were drawn (Fig. S5C-D). The gates were drawn in such a way that percent FRET positive cells (FRET %) (Fig. S5D) in the biosensor control group (treated with monomer) was always ≥ 1 as recommended in a prior study [77]. Upon confocal imaging of cells that expressed only acceptor (αSyn-mNG), there was no bleed-through of mNG into FRET channel (Fig. 3C) confirming the specificity of FRET-Flow cytometry. When rPFF was applied to the biosensor cells, αSyn aggregates were detected in FRET channel that were distinct from the background fluorescence of cells treated with αS monomer (Fig. 3D). αSyn-mTQ2 aggregates in biosensor cells were also detected in a different channel (Fig. 3D). Bleed-through from mTQ2 into FRET channel was eliminated by gating in FRET-Flow cytometer (Fig. S5D). When biosensor cells were treated with rPFF, 42.7% of single cells were found to be FRET positive suggesting αSyn oligomerization and aggregation events in this subset of cells (Fig. 3E). Integrated FRET median fluorescence intensity (MFI) was 19-fold higher (Fig. 3F) in rPFF-treated cells than those treated with αSyn monomer, suggesting significantly greater number of intracellular αSyn oligomerization and aggregation events taking place upon seeding with rPFF seeds. Overall, our results indicate that U251 αSyn biosensor cell line has been successfully generated and intracellular αSyn oligomerization and aggregation events are efficiently detected upon seeding by rPFF seeds.

**Figure 3:**
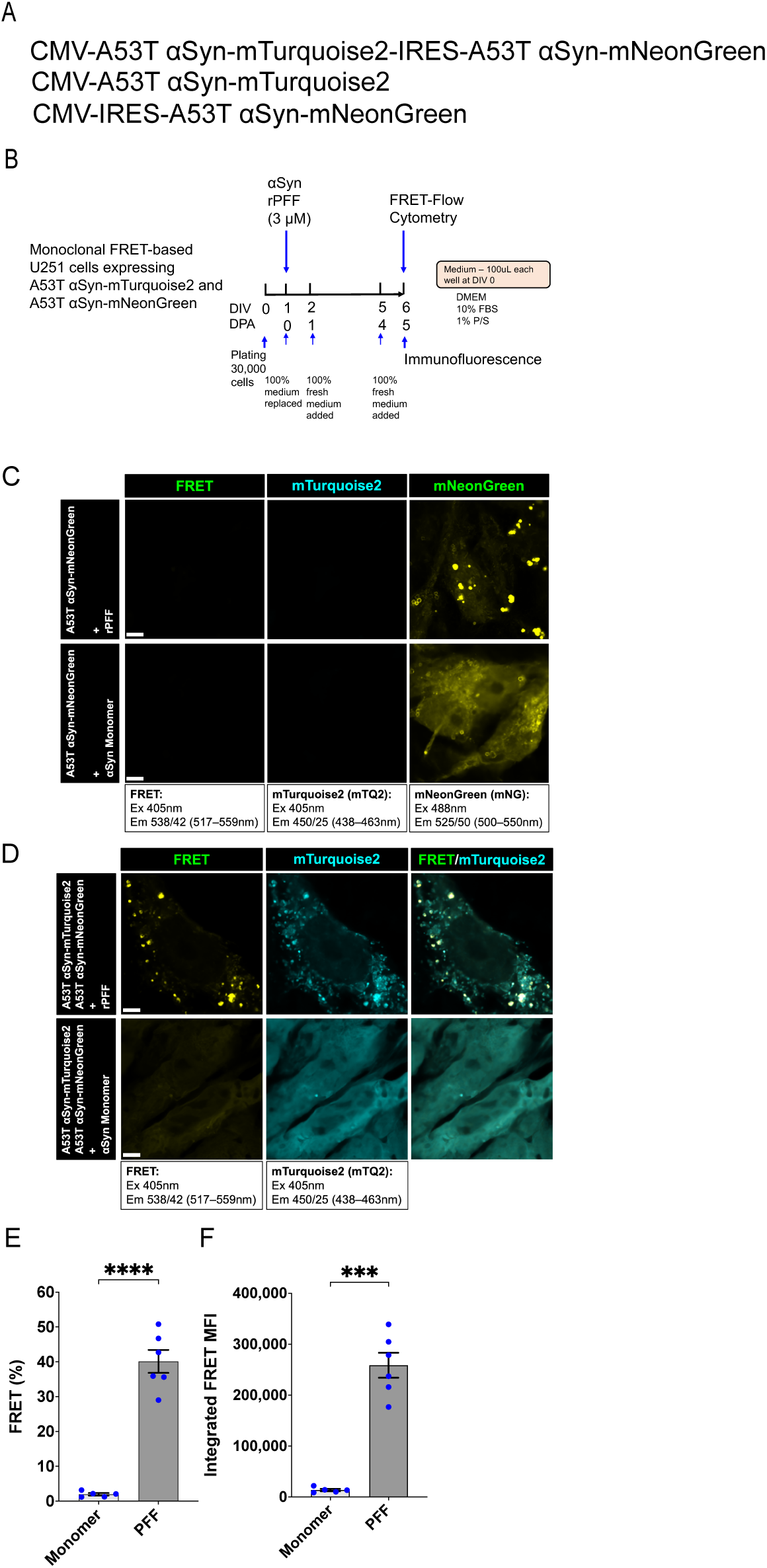
FRET-based αSyn biosensor U251 cells detect intracellular αSyn aggregation. (**A**) The lentiviral vectors for the expression of A53T αSyn-mTurquoise 2 and A53T αSyn-mNeonGreen (FRET pair) or A53T αSyn-mTurquoise 2 (FRET donor only control) or A53T αSyn-mNeonGreen (FRET acceptor only control) in U251 cells. (**B**) Experimental design showing the timelines of WT αSyn-rPFF application. Monoclonal biosensor cell line selection was performed using FRET-flow cytometry to select a cell line that has the highest naked seeding efficiency induced by αSyn rPFF. (**C, D**) Representative confocal images showing no bleedthrough of A53T αSyn-mNeonGreen into the FRET channel in FRET acceptor expressing cells and FRET signal in biosensor cells induced by rPFF. Scale bars: 5 µm. (**E, F**) Quantification of percent FRET positive cells and integrated FRET median fluorescence intensity (MFI) in αSyn rPFF and monomer treated biosensor cells by FRET-flow cytometry. Data are presented as mean ± SEM, and each dot corresponds to a biological replicate where 22,000 to 25,000 FRET biosensor single cells were analyzed; N=6 per group. Statistical analysis was performed by unpaired two-tailed student’s t-test with Welch’s correction. A p value <0.05 was considered significant. Significant differences are indicated by *p<0.05, **p<0.01, ***p<0.001, ****p<0.0001.

### Patient skin αSyn strains are amplified into amyloid fibrils by RT-QuIC

We previously reported that RT-QuIC assay amplifies and detects pathological αSyn seeds from the skin of PD, DLB, and MSA patients [48–50]. To investigate the pathology of different skin αSyn strains in U251 biosensor cells, we generated skin-derived αSyn amyloid fibrils (skin-amplified αSyn strains) from 2 cases each of PD, DLB, and MSA skin homogenates by RT-QuIC seed amplification assay. Similarly, 1 case each of PD, DLB, and MSA patient brain homogenate was used to generate brain-derived αSyn amyloid fibrils (brain-amplified αSyn strains) by RT-QuIC. Healthy skin and brain homogenates were used as controls to monitor the specificity of RT-QuIC assay. RT-QuIC run was completed when ThT fluorescence readout of skin and brain homogenates from healthy subjects reached a baseline set-point of 50,000 relative fluorescence units (RFU). Patient skin- and brain-derived αSyn strains resulted in increased ThT fluorescence readout ranging from 100,000 to 260,000 RFU within 29 hrs (Fig. 4A) and 56 hrs (Fig. 4B), respectively. These amplified skin and brain αSyn aggregates were subsequently purified by ultracentrifugation at 186,000xg to exclude monomeric αSyn. TEM analysis of purified skin and brain-amplified αSyn aggregates confirmed that they consist mostly of fibrillar (straight or twisted) or ribbon-like (flat) structures (Fig. 4C). A very small subpopulation of skin and brain-amplified PD and MSA αSyn fibrils exhibited twisted shape (Fig. 4C, black arrowheads). The DLB skin- and brain-amplified αSyn fibrils as well as *de novo* generated rPFF appeared to be mostly straight (Fig. 4C). Therefore, consistent with the previous reports [48–50], RT-QuIC assay amplifies αSyn strains from patient skin and brain homogenates into amyloid fibrils.

**Figure 4:**
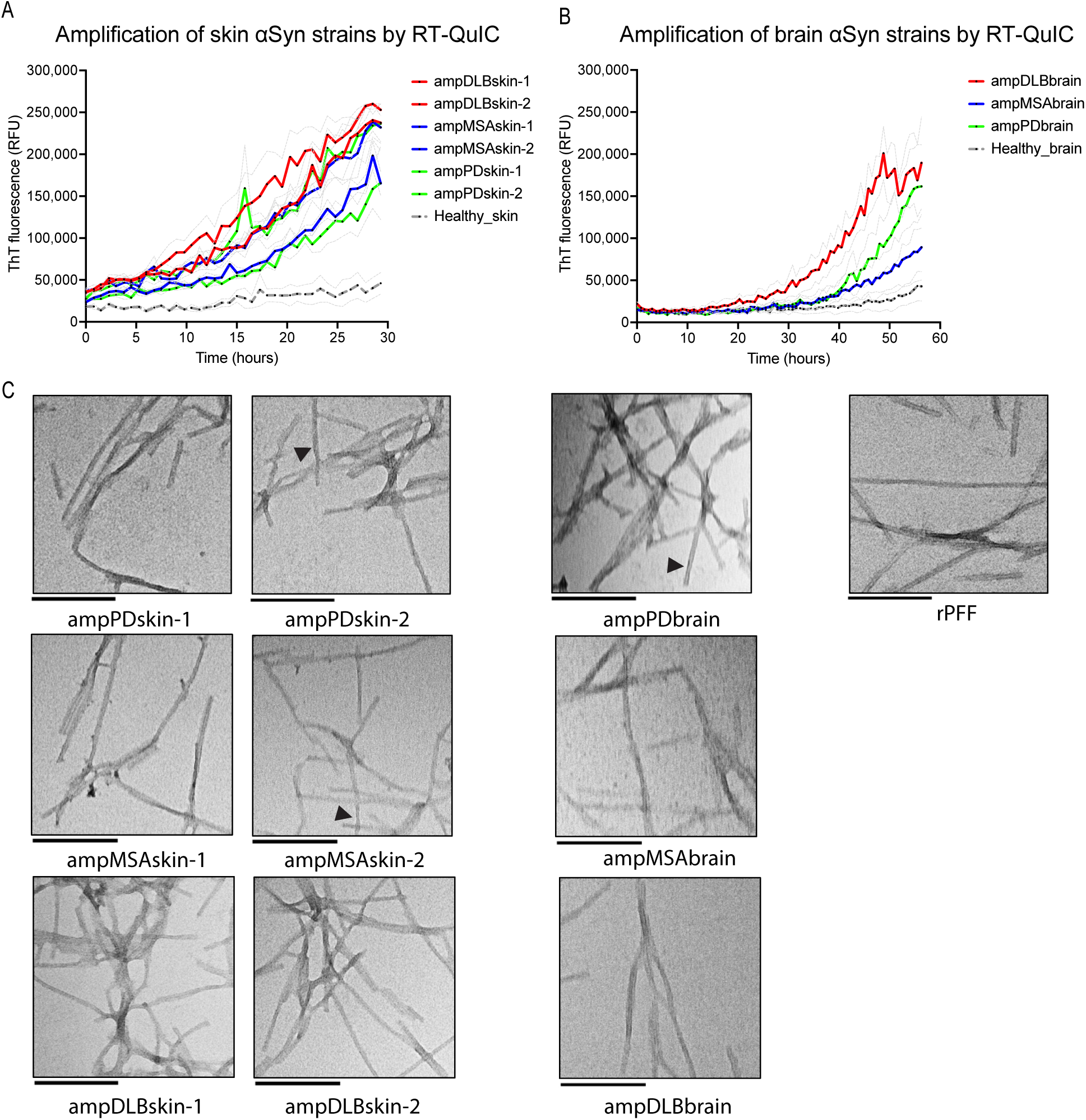
RT-QuIC seed amplification assay amplifies skin and brain-derived αSyn aggregates into amyloid fibrils. (**A, B**) Amplification of αSyn aggregates from PD, DLB, and MSA patient skin and brain homogenates using RT-QuIC. Data are presented as mean ± SEM (grey line), and each dot represents data analyzed from N=3 replicates. (**C**) Representative TEM images of PD, DLB, and MSA patient αSyn strains amplified from skin or brain homogenates. Black arrowheads indicate twists. Scale bars: 200 nm.

### Patient skin-amplified αSyn strains induce distinct inclusion pathology in U251 biosensor cells

Using U251 cells expressing A53T αSyn FRET biosensors, we sought to determine if patient skin-amplified PD, DLB, and MSA αSyn strains are pathologically active that can seed intracellular αSyn in biosensor cells. 100nM of purified PD, DLB, and MSA skin and brain-amplified αSyn strains were applied to U251 biosensor cells to seed intracellular aggregation of αSyn, along with rPFF and αSyn monomer as respectively positive and negative controls (Fig. 5A). Immunofluorescence of pathological p-αSyn inclusions and FRET-Flow cytometry was performed at day 4 post-treatments with patient-derived αSyn strains. Strong FRET signals were observed in U251 biosensor cells upon seeding by all patient-derived αSyn strains and rPFF but not the αSyn monomer control, and overlapped with the p-αSyn positive inclusions (Fig. 5B-E). The percent of biosensor cells positive for FRET (FRET %) was analyzed (Fig. S6A) and was normalized to the control monomer group to calculate FRET fold change as an indicator of seeding activity of a αSyn strain that induces intracellular FRET response. Remarkably, skin-amplified PD strains had higher seeding activity (FRET fold change) than skin-amplified DLB strains (Fig. 5F). There was no significant difference in the seeding activity between skin-amplified αSyn strains and brain-amplified αSyn strains from PD, DLB, and MSA patients (Fig. 5F). FRET efficiency, an indicator of aggregation propensity of a strain was calculated using integrated median fluorescence intensities of the FRET pairs. We found that FRET efficiency was also similar among patient skin and brain-amplified strains (Fig. S6B). Our results suggest that patient skin-amplified PD, DLB, and MSA αSyn strains are biologically active, and skin-amplified PD strains had a higher seeding activity than skin-amplified DLB strains in U251 biosensor cells.

**Figure 5:**
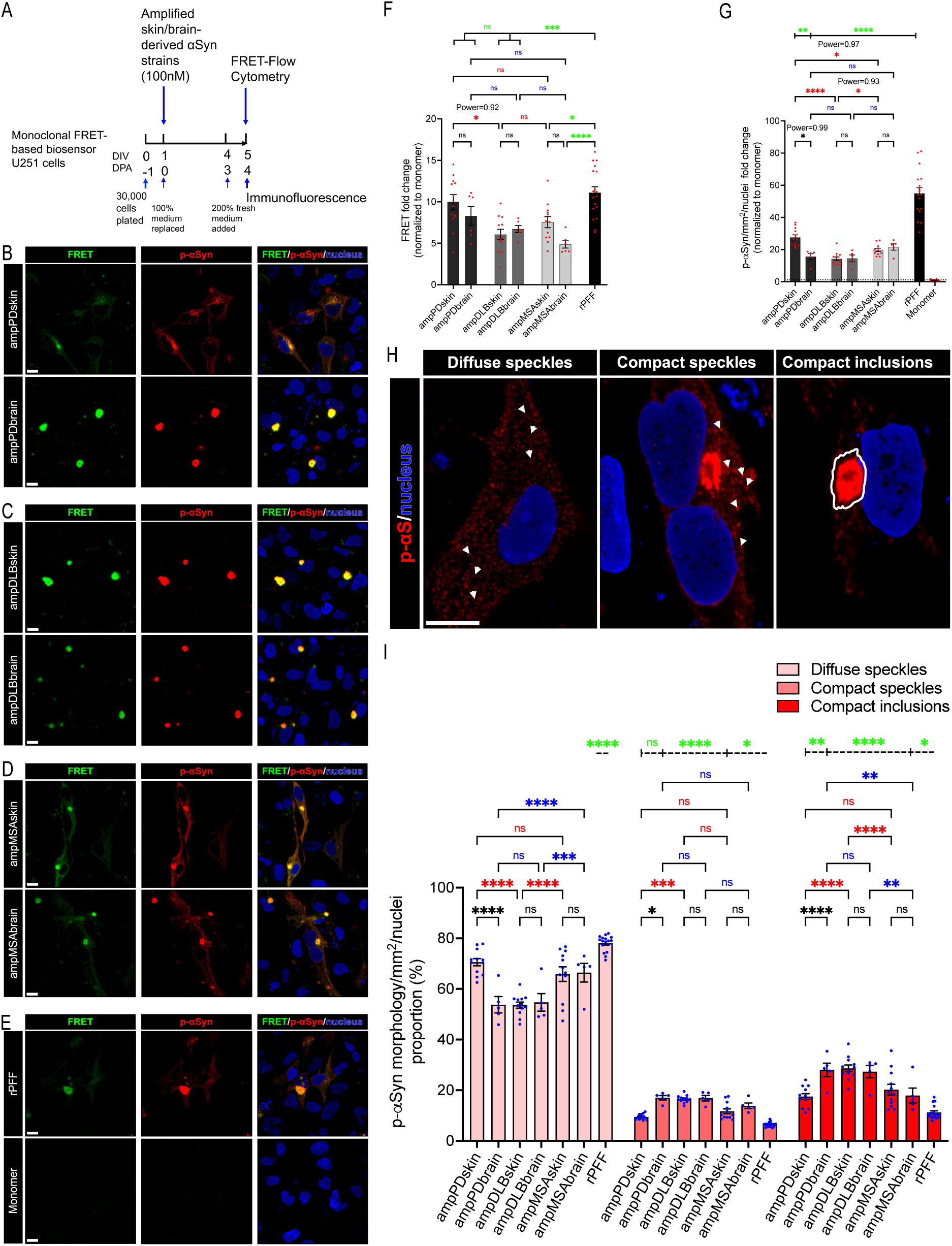
Patient skin-amplified αSyn strains induce distinct inclusion pathology in biosensor cells. (**A**) Experimental design showing the timelines of the application of skin or brain-amplified αSyn strains with lipofectamine on monoclonal FRET-based biosensor U251 cells. (**B-E, H**) Representative confocal images showing large FRET and p-αSyn positive Lewy body-like aggregates in the cytosol mostly tethered to the nuclear membrane, and classification (H) of p-αSyn aggregate morphology These inclusions were not present in αSyn monomer treated cells. Scale bars: 10 µm. (**F-G, I**) Quantification of FRET fold change (F), total intracellular p-αSyn aggregates (G), and proportions of p-αSyn aggregate morphology. Data are presented as mean ± SEM, and each dot corresponds to data analyzed from 20,000 to 25,000 (F) and 6,000 to 10,000 (G, I) FRET positive cells in a biological replicate from 2-3 independent experiments; N=12 (F) and N=10-12 (G, I) from the skin-amplified strains of 2 cases, N=6 (F) and N=5 (G, I) from the brain-amplified strains of 1 case, N=18 (F) and N=17 (G, I) from rPFF group, and N=18 (F) and N=9 (G, I) from αSyn monomer group. Statistical analysis was performed by ordinary one-way ANOVA with Tukey’s multiple comparison test or Brown-Forsythe and Welch ANOVA with Dunnet’s T3 multiple comparison test (F, G). A p value <0.05 was considered significant. Significant differences are indicated by *p<0.05, **p<0.01, ***p<0.001, ****p<0.0001.

To further examine morphological features of pathological intracellular αSyn aggregates in biosensor cells seeded by patient skin- and brain-amplified strains, immunofluorescence of p-αSyn inclusions in biosensor cells was performed at day 4 post-treatments with skin and brain-amplified strains and analyzed by high-content confocal microscope. FRET-positive inclusions were completely colocalized with pathological p-αSyn inclusions (Fig. 5B-E). Unexpectedly, skin-amplified PD strains induced significantly more p-αSyn inclusions than brain-amplified PD strains (Fig. 5G). Remarkably, skin-amplified PD strains induced significantly higher burdens of p-αSyn pathology than skin-amplified DLB and MSA strains (Fig. 5G). Moreover, skin-amplified MSA strains also induced more p-αSyn pathology than skin-amplified DLB strains (Fig. 5G). No significant differences were observed between skin- and brain-amplified strains in DLB and MSA groups in terms of their ability to induce p-αSyn positive inclusions (Fig. 5G). To enhance our understanding of p-αSyn inclusion morphology, we classified intracellular p-αSyn assemblies into 4 classes of morphologies based on mean fluorescence intensity (MFI) and size of the assembly, namely, diffuse speckles, diffuse inclusions, compact speckles, and compact inclusions (Fig. 5H). Lower MFI (≤ 14,999) was termed as diffuse, whereas higher MFI (≥15,000) was termed as compact. When a p-αSyn positive assembly covers a smaller area (0.5-4.5µm) in a cell, it was termed as a speckle, whereas, when it covers a larger area (4.6-35µm), it was termed as an inclusion. Using this classification, we found that skin- and brain-amplified αSyn strains induced more diffuse and less compact p-αSyn pathology overall (Fig. 5I). Remarkably, we found that skin-amplified PD strains induced more diffuse speckles and less compact inclusions than brain-amplified PD strains (Fig. 5I). PD Skin-amplified strains also differed from DLB-skin-amplified strains, the latter resulted in less diffuse speckles but more compact inclusions (Fig. 5I, red asterisks). In contrast, PD and DLB brain-amplified strains induced similar p-αSyn morphologies with more compact inclusions and less diffuse speckles than MSA brain-amplified strains (Fig. 5I, blue asterisks), fortifying the previously reported αSyn strain-specific differences between Lewy body diseases (PD or DLB) and MSA that used αSyn strains purified by sarkosyl from patient brain homogenate [43,78] or directly used patient brain homogenate [38,72]. DLB and MSA skin-amplified αSyn strains induced p-αSyn morphologies similarly as brain-amplified strains in U251 biosensor cells (Fig. 5I, black asterisks). The proportion of p-αSyn diffuse inclusions was very small and ranged from 1-4% that was not significantly different among groups. Taken together, our results indicate that PD, DLB and MSA patient-amplified αSyn strains recapitulate intracellular seeding activity and pathological p-αSyn inclusion morphology in U251 αSyn biosensor cells. Furthermore, skin-amplified αS strains from PD patients can be potentially discriminated from DLB patient skin-amplified αSyn strains by analyzing FRET seeding activity and p-αSyn inclusion morphology in U251 biosensor cells. Finally, distinct p-αSyn morphologies were induced by skin-amplified PD strains and brain-amplified PD strains, presumably suggesting the existence of distinct αSyn conformers in different tissues of PD patients.

### Patient skin-amplified αSyn strains trigger neurodegeneration

We have shown that patient skin-amplified αSyn strains are biologically active and mimic p-αSyn inclusion pathology of PD, DLB, and MSA in A53T αSyn expressing U251 biosensor cells. However, it is not known if such p-αSyn inclusion pathology is also recapitulated in human neurons. Therefore, we sought to reprogram human U251 glioblastoma biosensor cells into neurons while preserving the sensitive detection of αSyn aggregation. NEUROD1 has been reported to reprogram cultured human astrocytes in an Alzheimer’s disease mouse model into functional glutamatergic neurons [79]. Furthermore, U251 cells could be reprogrammed into functional glutamatergic neurons simply by NEUROD1 ectopic expression [63,80]. Here we used lentiviral vector to overexpress NEUROD1 in U251 glioblastoma biosensor cells using dsRed as a reporter (Fig. 6A). Indeed, NEUROD1 expressing U251 cells developed neuronal morphology with long neurites and were positive for neuronal marker TUJ1 and mature neuron marker NeuN at 48 DPI (Fig. 6B). Moreover, these mature neurons were VGLUT1 positive and negative for PROX1 (Fig. 6B) indicating that they are glutamatergic neurons but not granule cells. Untransduced biosensor cells surrounded TUJ1 positive neurons, potentially supporting neuronal maturation (Fig 6B). Patch-clamp generated action potentials in 80% of NEUROD1 positive cells as high sodium currents were observed on DPI 48 (Fig. 6C), suggesting successful reprogramming of U251 biosensor cells into human induced biosensor neurons (hiBNs). Skin and brain-amplified αSyn strains were applied to hiBNs on DPI 34 that resulted in the formation of intracellular pathological p-αSyn aggregates in TUJ1 positive (TUJ1+) neurons at DPI 48, which were mostly localized to the soma (cell body and the nucleus) of reprogrammed neurons (Fig. 6D-G). Quantification of pathological p-αSyn aggregates in hiBNs revealed similar seeding efficiency by skin- and brain-amplified strains (Fig. 6H). Length of p-αSyn aggregates was also measured by drawing the longest straight line within the aggregate, which revealed similar lengths of p-αSyn aggregates induced by skin and brain-amplified fibrils (Fig. 6I). In contrast, rPFF induced more p-αSyn aggregates that were longer than those induced by patient-derived αSyn strains. Finally, to assess neurotoxicity, number of surviving neurons were quantified by counting TUJ1+ soma at DPI 48, two weeks post-application of patient-amplified αSyn strains. We found that skin and brain-amplified strains were neurotoxic to 34-54% of total hiBNs as compared to the αSyn monomer-treated neurons, whereas rPFF was neurotoxic to 73% of total hiBNs (Fig. 6J). Brain-amplified PD fibrils were significantly more neurotoxic than the brain-amplified DLB fibrils (Fig. 6J). Furthermore, skin- and brain-amplified αSyn fibrils resulted in more dystrophic bulbous TUJ1+ neurites within 2 weeks in comparison to αSyn monomer control (Fig. 6D-G), indicating ongoing neurodegeneration of hiBNs induced by patient skin- and brain-amplified αSyn fibrils. In summary, these data show that patient skin- and brain-amplified αSyn strains can trigger the formation of pathological p-αSyn aggregates and neurodegeneration in human U251 biosensor cell-derived neurons.

**Figure 6:**
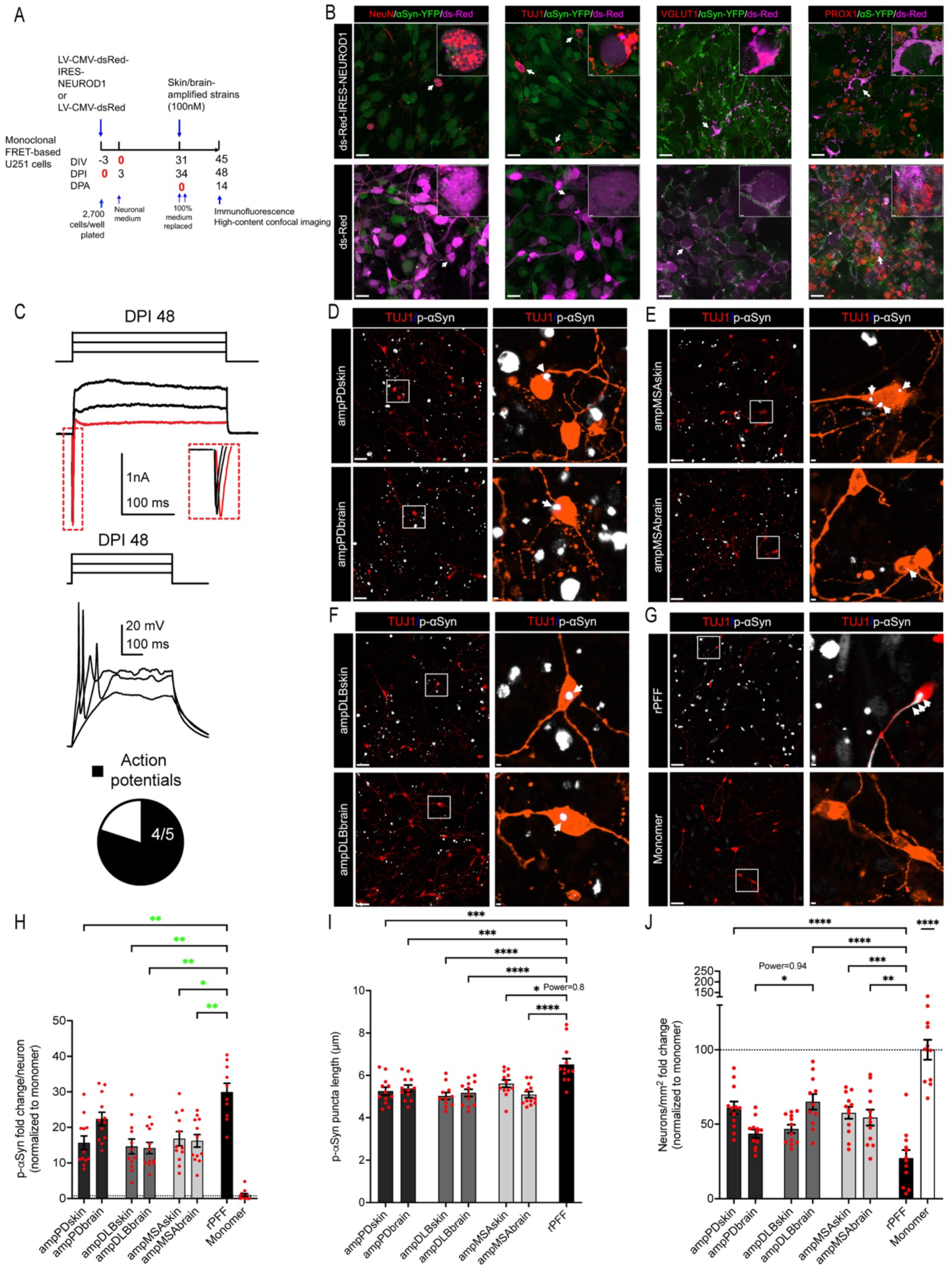
In αSyn biosensor induced neurons, skin- and brain-amplified αSyn fibrils trigger the formation of pathological p-αSyn inclusions and are neurotoxic. (**A**) Experimental design showing the timelines for generation of induced neurons from U251 FRET-based biosensor cells by reprogramming with lentiviral transduction of NEUROD1, and neuronal differentiation followed by application of skin- or brain-amplified αSyn strains with lipofectamine. (**B**) Representative confocal images showing the expression of TUJ1 and mature neuronal markers NeuN and VGLUT1, but not PROX1 at DPI 48. White arrows indicate the expanded inset image showing the subcellular localization of neuronal markers. Scale bars: 20 µm and 1µm. (**C**) A representative voltage clamp recording at -80 mV from a NEUROD1 positive cell at DPI 48 revealing large sodium currents in response to 50, 60 and 70 mV steps (300 ms); scale: 1nA/100ms. The sodium current in response to the 50 mV step is expanded and highlighted in red. Below, voltage traces from a NEUROD1 positive cell at DPI 48 in response to current injections (10, 30, 50 pA; 500 ms) revealing action potentials; scale 20 mV/100 ms. The pie chart highlights the number of NEUROD1 positive cells with action potentials. (**D-G**) Representative confocal images showing intraneuronal p-αSyn inclusions that were mostly localized in the soma. White squares indicate expanded image on the right showing the morphology and subcellular localization of p-αSyn inclusions as indicated by white arrows. Scale bars: 50µm and 3µm. (**H, I, J**) Quantification of p-αSyn inclusion fold change (H), p-αSyn inclusion lengths (I), and fold change of neuronal cell bodies (J) at DPA 14. Quantification of data are presented as mean ± SEM, and each dot corresponds to data analyzed from 180-700 neurons in a biological replicate. N=10-12 from the skin-amplified strains of 2 cases and N=11-12 from the brain-amplified strains of 1 case, N=10-12 from rPFF group, and N=11-12 from monomer group from 2 independent differentiations. Statistical analysis was performed by ordinary one-way ANOVA with Tukey’s multiple comparison test or Brown-Forsythe and Welch ANOVA with Dunnet’s T3 multiple comparison test (H). A p value <0.05 was considered significant. Significant differences are indicated by *p<0.05, **p<0.01, ***p<0.001, ****p<0.0001.

### Distinct p-αSyn signatures of patient skin αSyn strains dictate seeding activity *in vitro*

We have shown that patient skin-amplified αSyn strains can seed intracellular aggregation of αSyn into pathological Lewy body-like p-αSyn inclusions in biosensor cells (Fig. 5) and induced biosensor neurons (Fig. 6) that are reminiscent of Lewy body pathology found in PD, DLB, and MSA patient brains. However, it remains unknown if these newly formed intracellular αSyn fibrils generated in biosensor cells have intrinsic seeding activity to further propagate αSyn aggregation. To address this question, biosensor cells were treated with patient skin- and brain-amplified αSyn fibrils to seed intracellular aggregation of αSyn (Fig. 7A). When FRET positive αSyn inclusions were detected at day 4 post-application of patient-amplified αSyn strains, biosensor cells were lysed and sequentially purified by detergent extraction with NP-40 and sarkosyl (Fig. 7A). The detergent-insoluble fraction was then subjected to RT-QuIC assay and immunoblot. Detergent-insoluble fraction of PBS-treated biosensor cells (cellPBS) was found to be an ideal control as this group showed very minimal increase in ThT fluorescence that was set as a baseline (Fig. 7B). RT-QuIC reactions were stopped when cellPBS group reached baseline ThT fluorescence of 50,000 RFU (Fig. 7B). RT-QuIC assay detected significantly higher ThT fluorescence (hence higher *in vitro* seeding activity) from skin and brain αSyn strains derived from biosensor cells in comparison to control after 36 hours of RT-QuIC reactions (Fig. 7B). Comparison of *in vitro* RT-QuIC seeding activity at the endpoint 36 hour revealed no significant difference between biosensor cell-amplified skin and brain strains of DLB or MSA. Remarkably, skin PD strains exhibited higher RT-QuIC seeding activity than brain PD strains and skin DLB strains (Fig. 7C), which correspond to the higher burdens of intracellular p-αSyn inclusions triggered by PD skin-amplified αS strains in U251 biosensor cells as shown previously (Fig. 5H). On immunoblots, high p-αSyn levels were enriched in the detergent-insoluble fraction (Fig. 7E), unlike the detergent-soluble fraction (Fig. 7D), confirming that phosphorylated αSyn aggregates are detergent-insoluble. PD and MSA skin αSyn strains triggered higher levels of detergent-insoluble p-αSyn, unlike PD and MSA brain αSyn strains and DLB skin and brain αSyn strains (Fig. 7E). This potentially explains higher seeding activity of PD and MSA skin αSyn strains than DLB skin and brain αSyn strains and PD brain αS strains (Fig. 7C). Taken together, we showed that A53T αSyn U251 biosensor cells preserve seeding activity of PD, DLB, and MSA skin and brain αSyn strains which are faithfully propagated by *in vitro* RT-QuIC assay. The p-αSyn pathological burdens triggered in U251 biosensor cells dictate seeding activity of biosensor cell-derived patient skin and brain αSyn strains *in vitro*, allowing discrimination of skin αSyn strains between PD and DLB, and establishing a p-αSyn inclusion pathology to seeding relationship.

**Figure 7:**
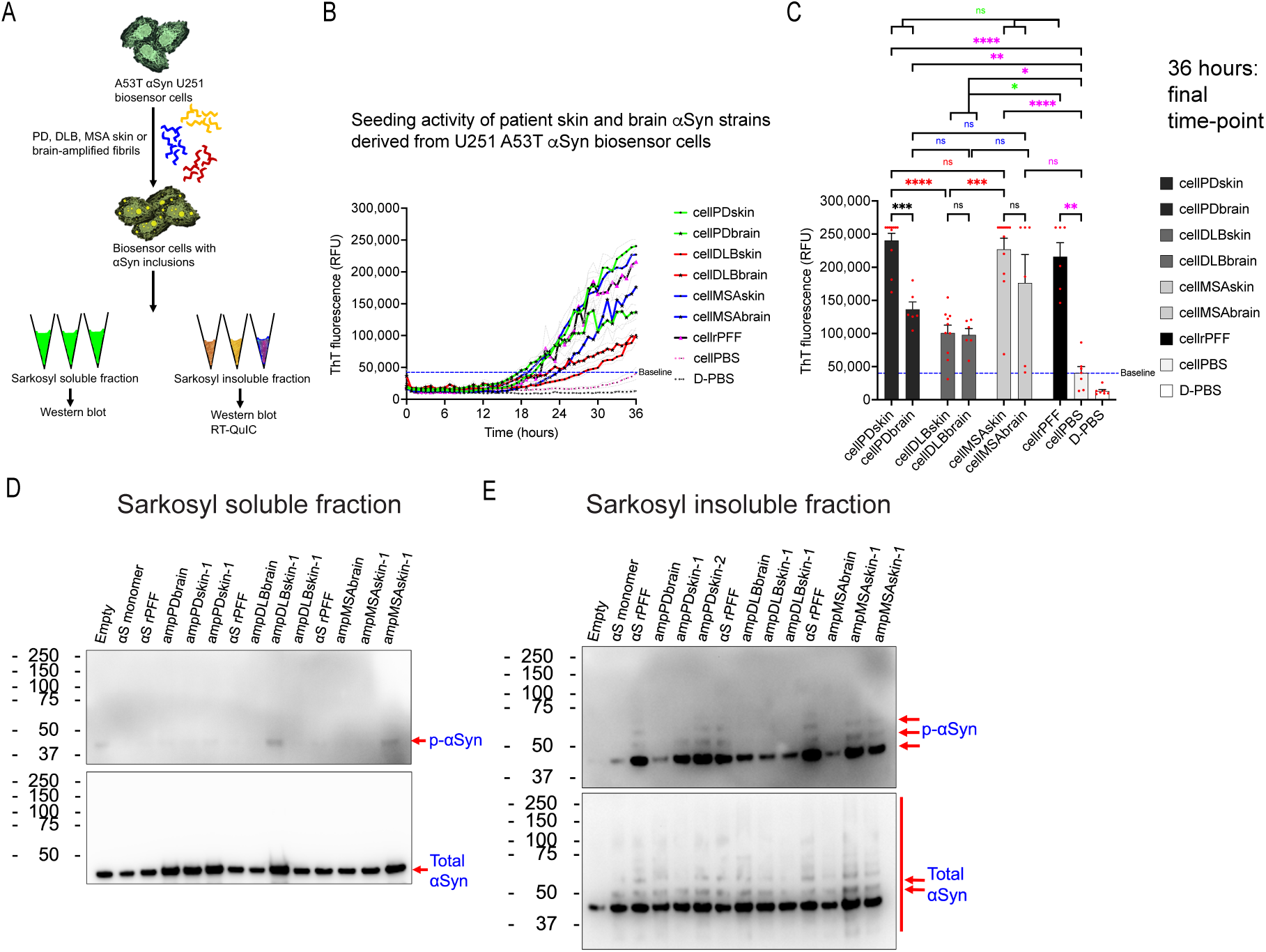
U251 biosensor cell-derived αSyn strains can be propagated *in vitro* by RT-QuIC. (**A**) Schematic of production and purification of U251 biosensor cell-derived αSyn strains after treatment with patient skin- or brain-amplified αSyn strains from cells. (**B**) RT-QuIC assay of biosensor cell-derived skin or brain αSyn strains showing their *in vitro* seeding activity. (**C**) Quantification of endpoint ThT fluorescence of RT-QuIC assay at 36-hour time-point showing *in vitro* seeding activity of patient skin- and brain-amplified αSyn strains following propagation in U251 biosensor cells. Data are presented as mean ± SEM. Each dot corresponds to data analyzed from N=10-12 replicates from biosensor cell-derived skin αSyn strains of 2 cases, N=6 replicates from biosensor cell-derived brain αSyn strains of 1 case, N=6, N=5, and N=7 replicates for biosensor cell-derived rPFF, αSyn monomer, and D-PBS, respectively, and N=7 replicates for RT-QuIC internal control with D-PBS quantified at every 45 minutes interval for 36 hours from 2 independent experiments. (**D-E**) Western blots showing pathological p-αSyn and total αSyn detected in sarkosyl detergent-soluble (D) and sarkosyl detergent-insoluble fraction (E) from U251 biosensor cells seeded with patient skin- or brain-amplified αSyn strains. Statistical analysis was performed by Brown-Forsythe and Welch ANOVA with Dunnet’s T3 multiple comparison test. A p value <0.05 was considered significant. Significant differences are indicated by *p<0.05, **p<0.01, ***p<0.001, ****p<0.0001.

## Discussion

It remains unknow how distinct αSyn strains are generated and propagated that give rise to distinct α-synucleinopathies such as PD, DLB, and MSA. We previously reported that pathological αSyn aggregates are present in PD, DLB, and MSA patient skin, and seeding activity of skin αSyn strains can be robustly detected by RT-QuIC assay *in vitro* [48–50]. However, disease-specific pathological features induced by PD, DLB, and MSA skin αSyn strains in a biological system remain unresolved.

In this study, we showed the existence of pathologically distinct αSyn strains from PD patient skin and brain as they trigger unique p-αSyn inclusion morphologies in U251 glioblastoma FRET-based αSyn biosensor cells. Further, we showed that DLB and MSA skin αSyn strains mimic p-αSyn inclusion morphologies of brain αSyn strains in biosensor cells. Moreover, skin αSyn strains induce degeneration of induced biosensor neurons like brain αSyn strains. Finally, we showed that the detergent-insoluble and pathological p-αSyn burdens in U251 biosensor cells dictates *in vitro* αSyn seeding activity with a potential to discriminate PD from DLB αSyn strains and thereby establishing a p-αSyn pathology to seeding relationship.

### U251 FRET-based αSyn biosensor cells for disease modeling

Many immortalized cell lines have been used to study αSyn aggregation [66,67]. Differentiation of SH-SY5Y human neuroblastoma cells to neuron-like cells have been used but they are difficult to grow and maintain, and widely differ in neuronal differentiation and functionality when compared in different laboratories, making them less reliable for αSyn aggregation studies [81]. By contrast, HEK293T cells have been widely used to study αSyn seeding by exogenous proteopathic seeds, inclusion morphology, and propagation [60,68]. Given that HEK293T cells were isolated from the kidney of a human embryo, the αSyn inclusions in HEK293T cells, induced by distinct αSyn strains, might not be a good representation of the inclusions originally formed in neurons, astrocytes or oligodendrocytes in the human brain, because it has been shown that intracellular milieu generates distinct pathological strains of αSyn [46]. Therefore, we used human glioblastoma cell lines that grow rapidly and are easy to maintain. As prior work showed that exogenous αSyn rPFF can seed intracellular aggregation of αSyn to form inclusions [27,28,75], we expressed A53T αSyn-EGFP in different glioblastoma cell lines and exposed them to exogenous αS rPFF. Upon screening, we found that human U251 cells can efficiently seed intracellular A53T αSyn-EGFP into large Lewy body-like detergent-insoluble inclusions (Fig. 1C). We subsequently generated monoclonal U251 cells stably expressing A53T or WT αSyn-EGFP. When these cells were exposed to exogenous rPFF, intracellular detergent-insoluble p-αSyn inclusions were readily detected, suggesting efficient seeding by rPFF and formation of pathological A53T or WT p-αSyn inclusions in U251 cells (Fig. 2C). To our knowledge, rapid *in situ* detection of intracellular detergent-insoluble αSyn inclusion in cells has not been shown by previous studies with existing cellular models of αSyn aggregation. Strikingly, we found that U251 cells allowed rapid detection of bright EGFP positive intracellular αSyn inclusions following a simple treatment with a non-ionic detergent (Fig. S1A), suggesting that U251 cells can be used for powerful analyses of detergent-insoluble αSyn aggregates inside cells with tools like high-content imaging. To track αSyn inclusion formation and morphology in live cells, a FRET-based biosensor cell model using human U251 biosensor cells was generated. To express equal proportion of donor and acceptor fluorophores, a bicistronic lentiviral vector was designed that allowed the expression of fusion proteins A53T αSyn-mTurquoise2 (αSyn-mTQ2 or αSyn-CFP) controlled by CMV promoter and A53T αSyn-mNeonGreen (αSyn-mNG or αSyn-YFP) controlled by IRES (Fig. 3A). Using FRET-Flow cytometry, single-cell population of U251 biosensor cells was gated to report percent of cells that were FRET positive (FRET %) (Fig. S5). We did not detect any bleed through of αSyn-mNG acceptor into FRET channel either by FRET-Flow cytometry or confocal imaging in U251 biosensor cells (Fig. 3C, S5C), confirming an ideal choice of donor and acceptor fluorophores. Intracellular αSyn inclusions in single-cells were efficiently detected by FRET-Flow cytometry or confocal imaging when rPFF seeds were applied to U251 biosensor cells (Fig. 3D, S5D). This newly generated monoclonal FRET-based αSyn U251 cell model provides a robust platform to investigate intracellular αSyn inclusion pathology induced by patient-derived αSyn strains.

### PD, DLB, and MSA skin αSyn strains: inclusion pathology to seeding relationship

Here, we showed that skin-derived αS strains from PD, DLB, and MSA patients impart distinct biological seeding activity and inclusion pathology in human U251 biosensor cells. When patient skin αSyn strains were amplified by RT-QuIC, we observed a mixture of mostly straight and a small subset of twisted filaments in PD and MSA skin- and brain-amplified fibrillary structures using TEM (Fig. 4C) which is also shown in prior work reported by others [3,40,42,78,82]. Mostly, straight filaments were observed in DLB skin- and brain-amplified assemblies (Fig. 4C). Perren et al. showed that PMCA-amplified PD and MSA αSyn strains shared similar profile of partial proteinase-K digestion resembling ribbon-like morphology, which was different than that of PMCA-amplified DLB αSyn strains resembling fibril-like morphology [40] consistent with our findings in the context of TEM imaging. In contrast, Tanudjojo et al. observe that PMCA-amplified MSA αSyn strains resembled fibril-like morphology after partial proteinase-K digestion which was different from ribbon-like morphology of PMCA-amplified PD αSyn strains [42]. Furthermore, some studies show that αSyn strains amplified from brain homogenates are structurally different from αSyn strains present in brain homogenates [82,83]. Notably, these studies have used different salt concentrations or different buffer and at different pH and temperatures to amplify αS strains that has been shown to affect *de novo* generated αSyn fibril conformation [35], which might possibly be one of the reasons for structural differences between these preparations. On the other hand, isolation and purification of αSyn fibrils from patient brain might potentially induce conformational changes in patient αSyn strains [84]. Therefore, studies involving patient-derived αSyn assemblies require careful investigation and the availability of robust biosensor cells may permit discrimination of different αSyn strains under quasi-physiological conditions.

Interestingly, we found that seeding activity (FRET fold change) of skin-amplified PD strains in biosensor cells is higher than that of skin-amplified DLB strains (Fig. 5F). Further, we showed that skin-amplified PD strains induce greater number of pathological p-αSyn inclusions than skin-amplified DLB strains in biosensor cells (Fig. 5G), which is consistent with FRET-based seeding activity. Conversely, lesser p-αSyn inclusions were induced by brain-amplified PD strains than skin-amplified PD strains. To further enhance our understanding, we thoroughly classified pathological p-αSyn inclusion morphology induced by skin and brain-amplified αSyn strains in U251 biosensor cells using high-content confocal imaging. Skin- and brain-amplified groups of DLB and MSA induced similar p-αSyn inclusion morphology. Moreover, rPFF-induced p-αSyn morphologies are distinct from patient skin and brain-amplified strains which is consistent with recent work [44,45]. Remarkably, we found that PD and MSA skin-amplified αSyn strains impart similar p-αSyn inclusion morphologies that were distinct from skin-amplified DLB strains in U251 biosensor cells (Fig. 5I). Even though to the best of our knowledge, we are the first to report the pathology of patient skin-amplified strains in cultured cells, the outcome of our findings were found to be consistent with a previous study where PMCA-amplified PD αSyn strains from brain homogenates induced p-αSyn pathology in a greater number of cells than PMCA-amplified DLB αSyn strains *in vivo* [40]. In contrast, PD and DLB brain-amplified αS strains impart similar p-αSyn inclusion morphologies that were distinct from brain-amplified MSA strains in U251 biosensor cells (Fig. 5I). This was found to be consistent with prior studies that use patient brain homogenates in HEK293T cells or RT-QuIC-amplified αSyn strains from patient CSF [38,72,85]. In summary, our findings were consistent with prior studies that have used patient αSyn strains either directly from brain homogenates [37,38,41,72] or PMCA-amplified αSyn strains from patient brain homogenates [42,45] or both [40,44] to study pathological features of patient αSyn strains in cultured cells or in mice. It is noted that a prior work reported that PD brain homogenate-derived αSyn strains cannot seed A53T αSyn-YFP expressed in HEK293T cells [37] However, we showed here that skin-amplified PD αSyn strains triggered αSyn seeding and formation of detergent-insoluble and pathological p-αSyn inclusions in glioblastoma U251 biosensor cells expressing A53T αS-mTQ2 and A53T αS-mNG concurrently. Our findings are supported by previous work that detected αSyn seeding from PD brain homogenates in HEK293T biosensor cells expressing A53T αSyn-CFP and A53T αSyn-YFP [38]. Remarkably, our results showed that distinct p-αSyn morphology was induced by skin-amplified and brain-amplified PD strains (Fig. 5I), supporting the existence of potentially distinct αSyn conformers in PD patients.

An additional advantage of U251 biosensor cells is that they can be directly reprogrammed into neurons using NEUROD1 transcription factor as shown previously [63], allowing for the investigation of intracellular aggregation of αSyn in human induced biosensor neurons (hiBNs) from the same cell lineage. Unlike prior studies [63], we showed that hiBNs are physiologically functional when matured by day 48 and can generate action potentials upon current injections followed by electrophysiological recordings (Fig. 6B-C). Upon further characterization of our hiBNs, we found that hiBNs are glutamatergic neurons that lack PROX1 indicating that hiBNs are not granule cells. hiBNs morphologically resembled CA3 pyramidal neurons where the apical dendrite branches closer to the soma as shown in a prior study [86]. Skin and brain-amplified αSyn strains and rPFF induced round or oval p-αSyn inclusions that were localized in the soma of hiBNs (Fig. 6D-G) which is consistent with the prior work describing the localization of αSyn inclusions in PD, DLB, and MSA brains [1,3] and studies involving αSyn inclusions in the hippocampus [2,4,87,88]. Skin and brain-amplified strains were found to be neurotoxic that resulted in bulbous dystrophic neurites (Fig. 6J) as previously reported while using patient PD and MSA-amplified fibrils [42]. These results suggest that skin and brain-amplified strains trigger neurodegeneration.

An open question is whether the nascent αSyn aggregates generated in U251 biosensors in response to seeding by patient-derived αSyn strains retain strain-specific properties. We thus performed RT-QuIC assay using detergent-insoluble fraction purified from FRET positive U251 biosensor cells treated by skin and brain-amplified strains. Strikingly, we found that *in vitro* RT-QuIC seeding activity (Fig. 7C) of skin-amplified PD strains is distinct from brain-amplified PD strains and skin-amplified DLB strains (Fig. 7C). This pattern is remarkably preserved in terms of the FRET-based seeding potential (Fig. 5F) and the p-αSyn burden (Fig. 5G). This result fortified our finding that possibly distinct αSyn conformers are present in PD patients. Further, RT-QuIC assay of U251 biosensor cell-derived skin αS strains allows us to potentially discriminate PD and DLB strains using the patient skin.

In summary, we show the existence of pathologically distinct αSyn strains from PD patient brain and skin as revealed by U251 biosensor cells. Further, patient skin-amplified αSyn strains mimic pathological features of PD, DLB, and MSA in U251 biosensor cells and cause degeneration of biosensor cell-derived induced neurons. Finally, we show that the pathological αSyn inclusion burdens triggered by skin αSyn strains in U251 biosensor cells dictate their seeding activity in cells and *in vitro* as revealed by FRET-Flow cytometry and RT-QuIC assay respectively, thus establishing αSyn pathology to seeding relationship.

### Advantages of U251 αSyn biosensor cells

U251 αSyn biosensor cells are a powerful model to interrogate disease-specific αSyn strains because these rapidly dividing cells can internalize αSyn seeds and trigger intracellular αSyn aggregation, can be used to easily detect detergent-insoluble αSyn aggregates just within 3 days of exogenous αSyn seeding, can be directly reprogrammed to postmitotic electrophysiologically functional neurons by a single transcription factor, and can potentially discriminate PD and DLB skin-derived αSyn strains. U251 biosensor cell-model thus can be used for studying pathophysiological mechanisms that control PD, DLB, or MSA strain-specific aggregation and receptor-mediated internalization, unraveling novel synucleinopathy-specific biomarkers, and facilitating high-throughput screening of small molecules or biologics that might prevent or alleviate intracellular αSyn aggregation leading to potential disease-modifying therapeutic interventions.

## Conclusions

We find that PD skin-derived αSyn strain is pathologically distinct from PD brain-derived αSyn strain because it triggers a unique phenotype of pathological p-αSyn inclusions in U251 αSyn biosensor cells, suggesting the existence of distinct αSyn conformers from different tissues in PD patients. The DLB and MSA patient-derived αSyn strains are also competent to induce intracellular p-αSyn inclusions in these biosensor cells. We have demonstrated a consistent relationship between intracellular p-αSyn pathological burden and seeding activity induced by patient-derived αSyn strains in U251 biosensor cells. Importantly, U251 FRET-based αSyn biosensor cell model is a robust tool to potentially discriminate αSyn strains in PD and DLB and study disease-specific etiology and pathogenesis using a variety of patient-derived tissues.

## List of abbreviations

αSyn: alpha-synuclein
p-αSyn: phosphorylated αSyn at serine 129
PD: Parkinson’s disease
DLB: Dementia with Lewy bodies
MSA: Multiple system atrophy
rPFF: recombinant alpha-synuclein pre-formed fibrils
RT-QuIC: Real-time quaking-induced conversion
PMCA: Protein misfolding cyclic amplification
FRET: Förster (or Fluorescence) resonance energy transfer
MFI: Median/Mean fluorescence intensity
NEUROD1: Neuronal Differentiation 1
hiBNs: human induced biosensor neurons
ThT: Thioflavin T
WT: Wild type
LV: Lentivirus

## Declarations

### Ethics approval

Not applicable.

### Consent for publication

The authors give consent for publication.

### Availability of data and materials

All data are available in the main text or supplementary material.

### Authors’ contribution

A.R. conceived the study, designed and performed most of the experiments (except as noted below), constructed lentiviral vectors, generated monoclonal U251-αSyn FRET-based stable biosensor cell line, developed protocols for automated image acquisition and quantification on high-content confocal imaging system, acquired, analyzed, and interpreted the data, performed statistical analyses, prepared figures and graphical abstract, and wrote the draft manuscript. W.W. provided administrative and technical support. J.C.G. and X.Y. acquired, analyzed, and interpreted the data from electrophysiology of induced neurons. L.O.W. and J.I.W. interpreted the data and supervised the experiment associated with electrophysiology. C.-L.Z. and S.G.C. supervised and coordinated the overall study, revised the manuscript, and obtained funding.

## Acknowledgements

We are grateful to Connor Bargar for sharing his technical experience with RT-QuIC assay. We thank UAB High Resolution Imaging Facility for supporting this study. Lastly, we want to acknowledge the UAB Flow Cytometry and Single Cell Core facility supported by the NIH AI027767 (Center for AIDS Research) and CA013148 (The O’Neal Comprehensive Cancer Center) grants. We also acknowledge the use of the de-identified tissue samples obtained through the NIH NeuroBioBank.

---

**Figure S1:**
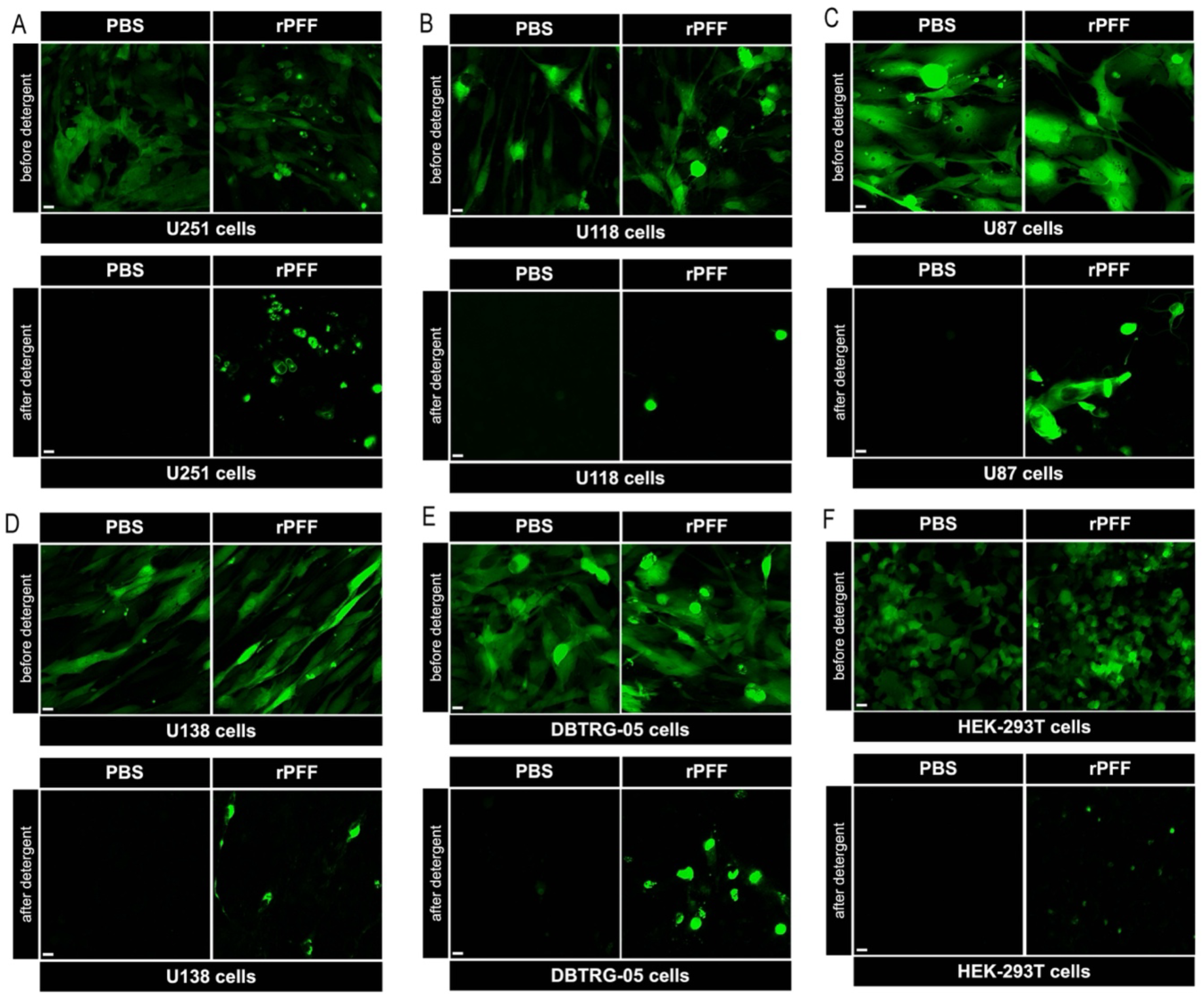
Application of detergent on live glioblastoma cells reveals detergent-insoluble intracellular αSyn-EGFP aggregates. (**A-F**) Representative live images before and after detergent treatment reveals total αSyn-EGFP and detergent-insoluble αSyn-EGFP inclusions in the indicated cell lines after application of WT αSyn rPFF or D-PBS at DPA 3 (DIV/DPI 7). Scale bars: 10µm. DIV: days *in vitro*, DPA: days post-application of rPFF, DPI: days post-infection with lentivirus.

**Figure S2:**
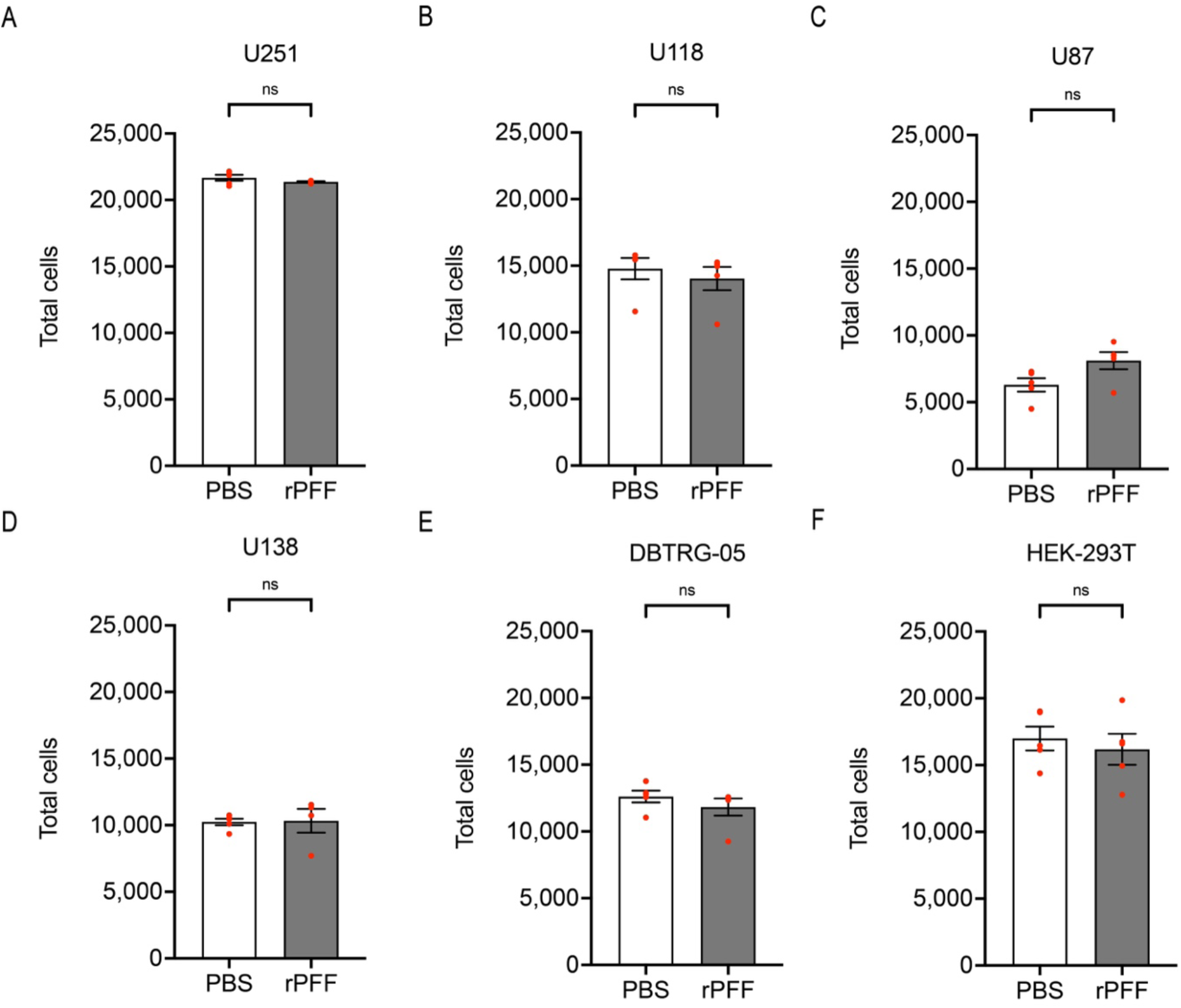
Total glioblastoma cells counted by high-content imaging. (**A-F**) Quantification of total cells positive for DAPI following PBS and rPFF treatments in the indicated cell lines at DPA 3 (DIV/DPI 7). Data are presented as mean ± SEM and each dot corresponds to a biological replicate; N=4-5 per group. Statistical analysis was performed by unpaired two-tailed Student’s t-test with (A, D) or without Welch’s correction (C, F) or unpaired two-tailed Mann-Whitney test (B, E) and shown as *p<0.05, **p<0.01, ***p<0.001, ****p<0.0001.

**Figure S3:**
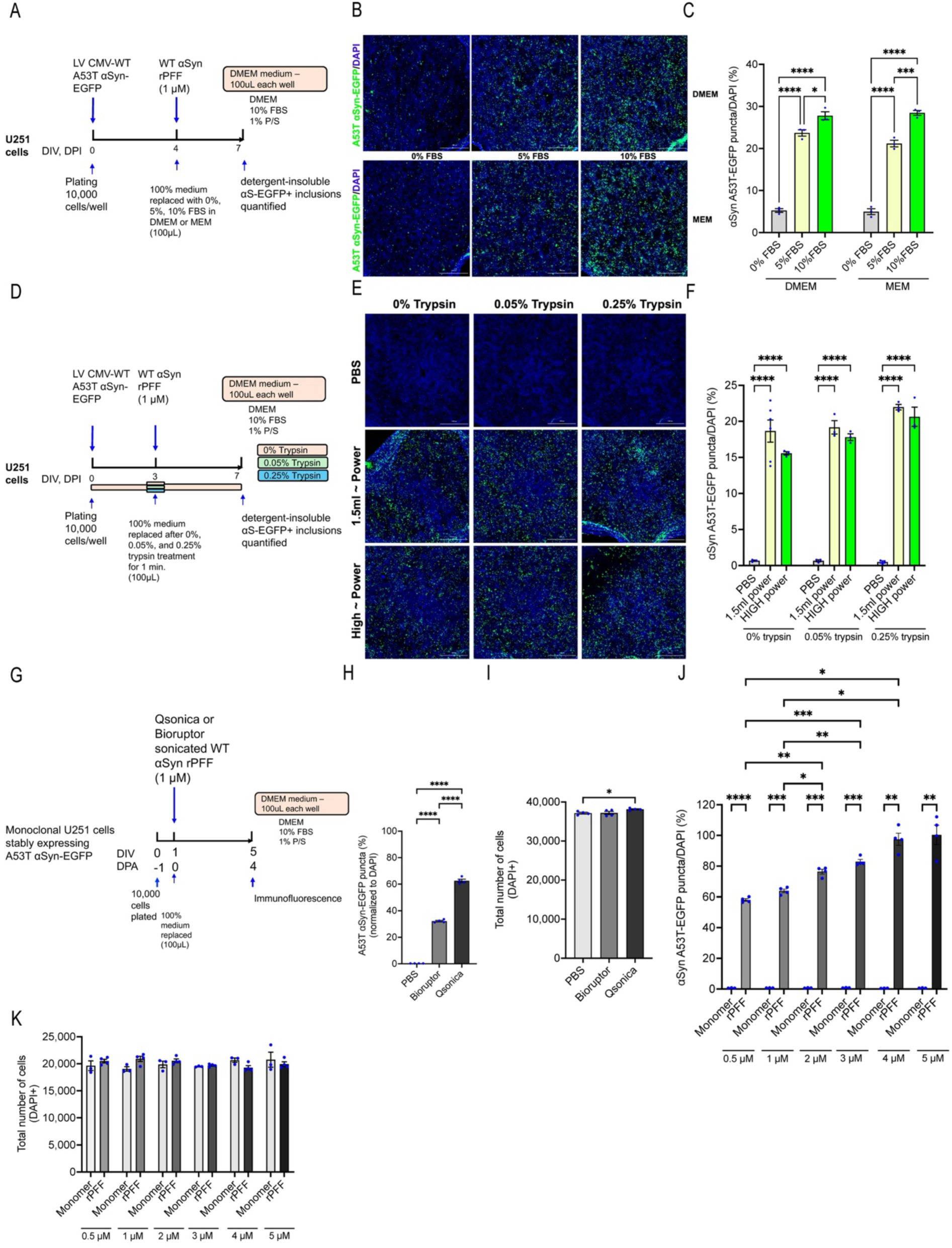
Optimizing U251 cell culture conditions for enhancing naked seeding with αSyn rPFF. (**A, D, G**) Experimental design showing the timelines of lentivirus and αSyn rPFF application with medium changes and trypsin pre-treatment. (**B, E**) Representative images showing detergent-insoluble A53T αSyn-EGFP inclusions (green) and cell nuclei (DAPI) in the indicated cell culture conditions. (**C, F, H, I, J, K**) Quantification of detergent-insoluble A53T αSyn-EGFP inclusions and total cells for the indicated cell culture and αSyn rPFF sonication conditions. Data are presented as mean ± SEM, and each dot corresponds to data analyzed from 20,000-36,000 cells on average in a biological replicate; N=3-5 per group. Statistical analysis was performed by ordinary one-way ANOVA with Tukey’s multiple comparison post-hoc test or by Brown-Forsythe and Welch ANOVA with Dunnet’s T3 multiple comparison test (I, J) and shown as *p<0.05, **p<0.01, ***p<0.001, ****p<0.0001.

**Figure S4:**
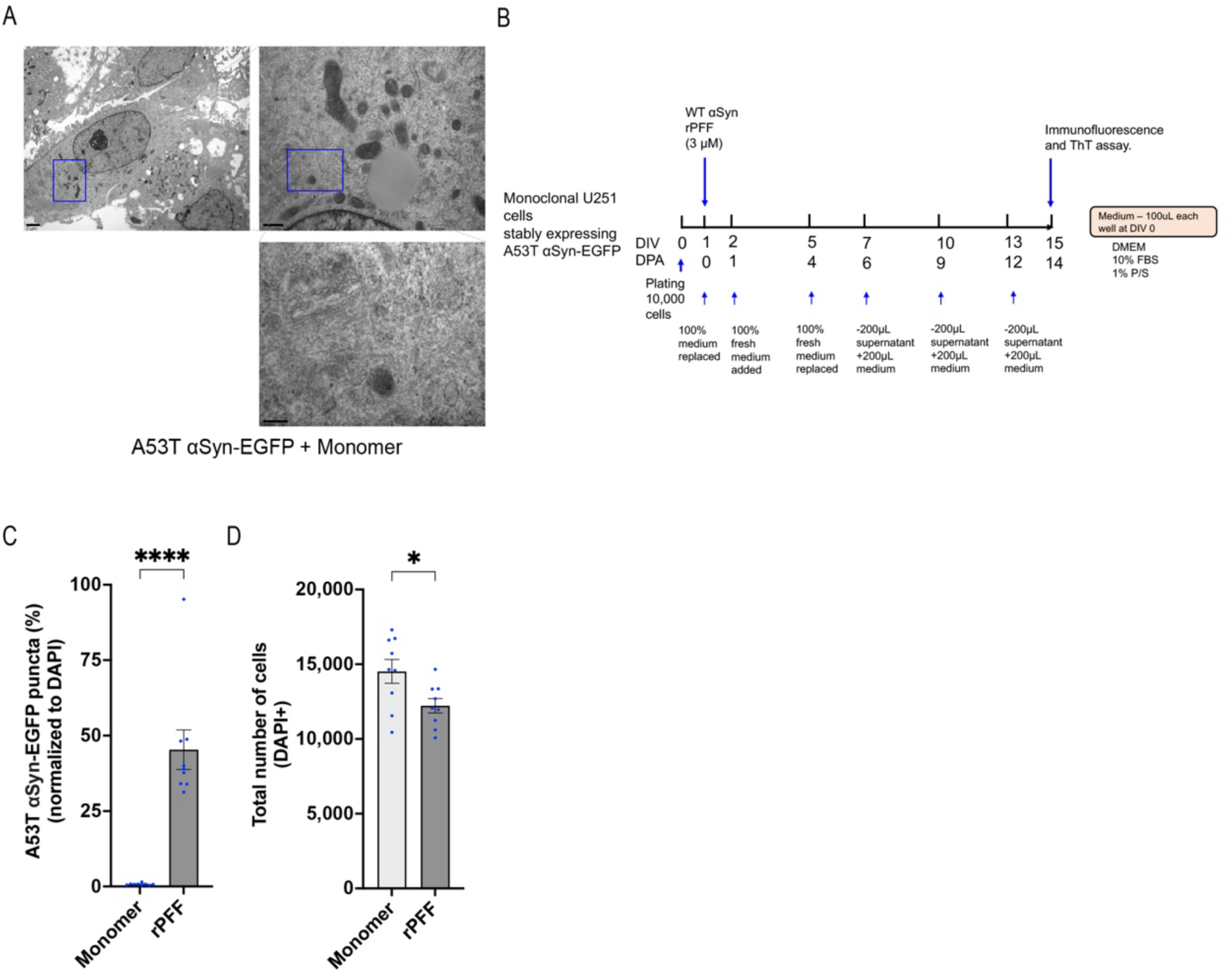
αSyn monomer cannot induce intracellular αSyn aggregation unlike rPFF in a long-term culture of U251 cells. (**A**) Representative TEM images of αSyn monomer treated U251 cells expressing A53T αSyn-EGFP showing the absence of fibrillar-like aggregate structures. Scale bars: 2 µm (3,200x), 0.5 µm (16,000x), and 0.2 µm (50,000x). (**B**) Experimental design showing the timelines rPFF or αSyn monomer application and medium changes. (**C, D**) Quantification of A53T αSyn-EGFP inclusions and total number of DAPI positive cells at DPA14 (DIV15) of αSyn rPFF or monomer treatment. Data are presented as mean ± SEM, and each dot corresponds to data analyzed from 10,000 to 15,000 cells in a biological replicate; N=9-10 per group. Statistical analysis was performed by unpaired two-tailed Mann Whitney test or unpaired two-tailed student’s t-test. A p-value <0.05 was considered significant. Significant differences are indicated by *p<0.05, **p<0.01, ***p<0.001, ****p<0.0001.

**Figure S5:**
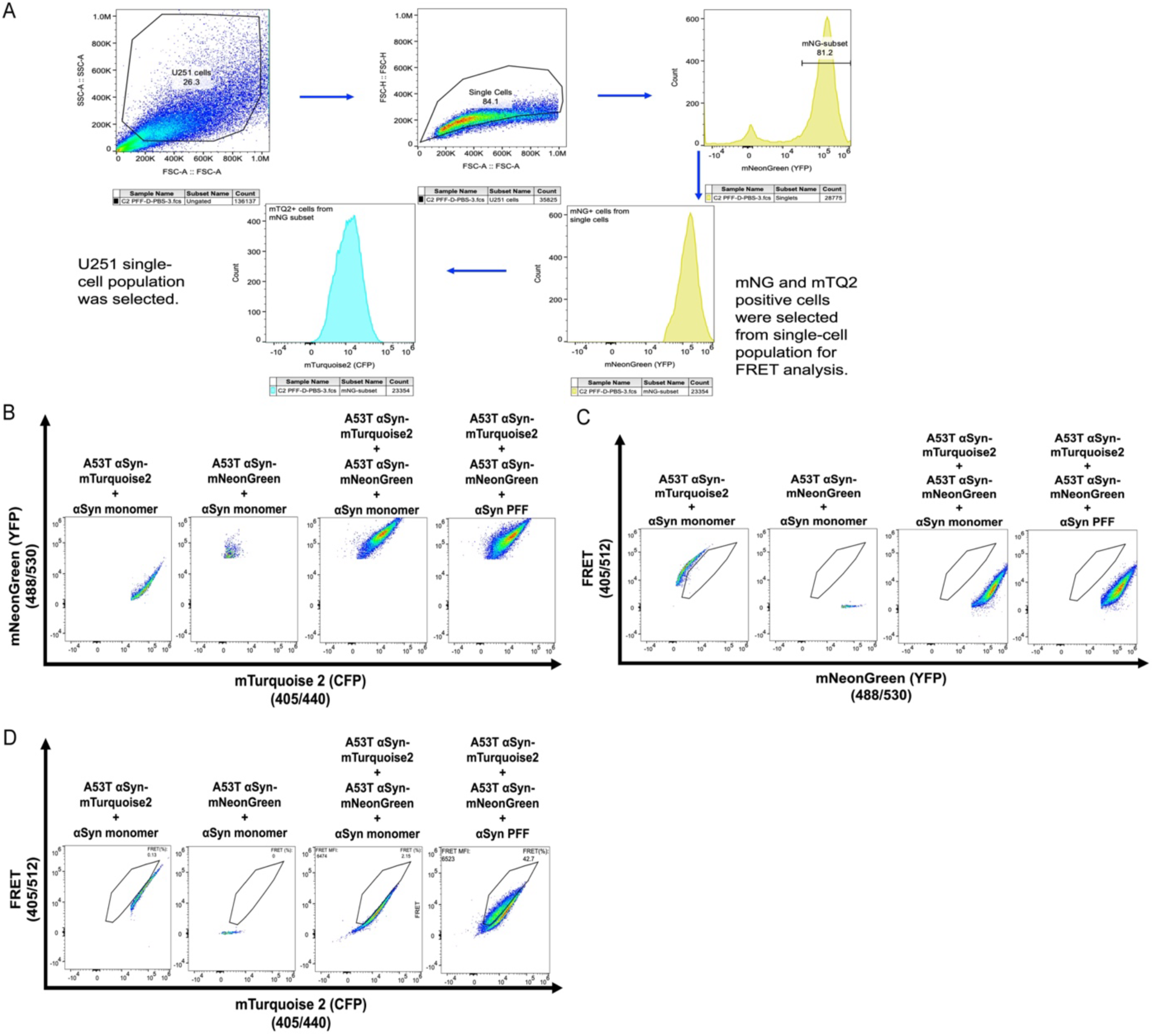
FRET gating strategy to quantify percent FRET positive cells using FRET-Flow cytometry. (**A**) Representative plots with gating to detect single U251 biosensor cells that express both αSyn-mTurquoise2 and αSyn mNeonGreen using FRET-flow cytometry. (**B, C, D**) Representative plots with appropriately designed gates to detect FRET with the indicated combination of lasers (excitation/emission) showing the cellular expression profile of mTurquoise2 or mNeonGreen or FRET with either αSyn monomer or rPFF treatment.

**Figure S6:**
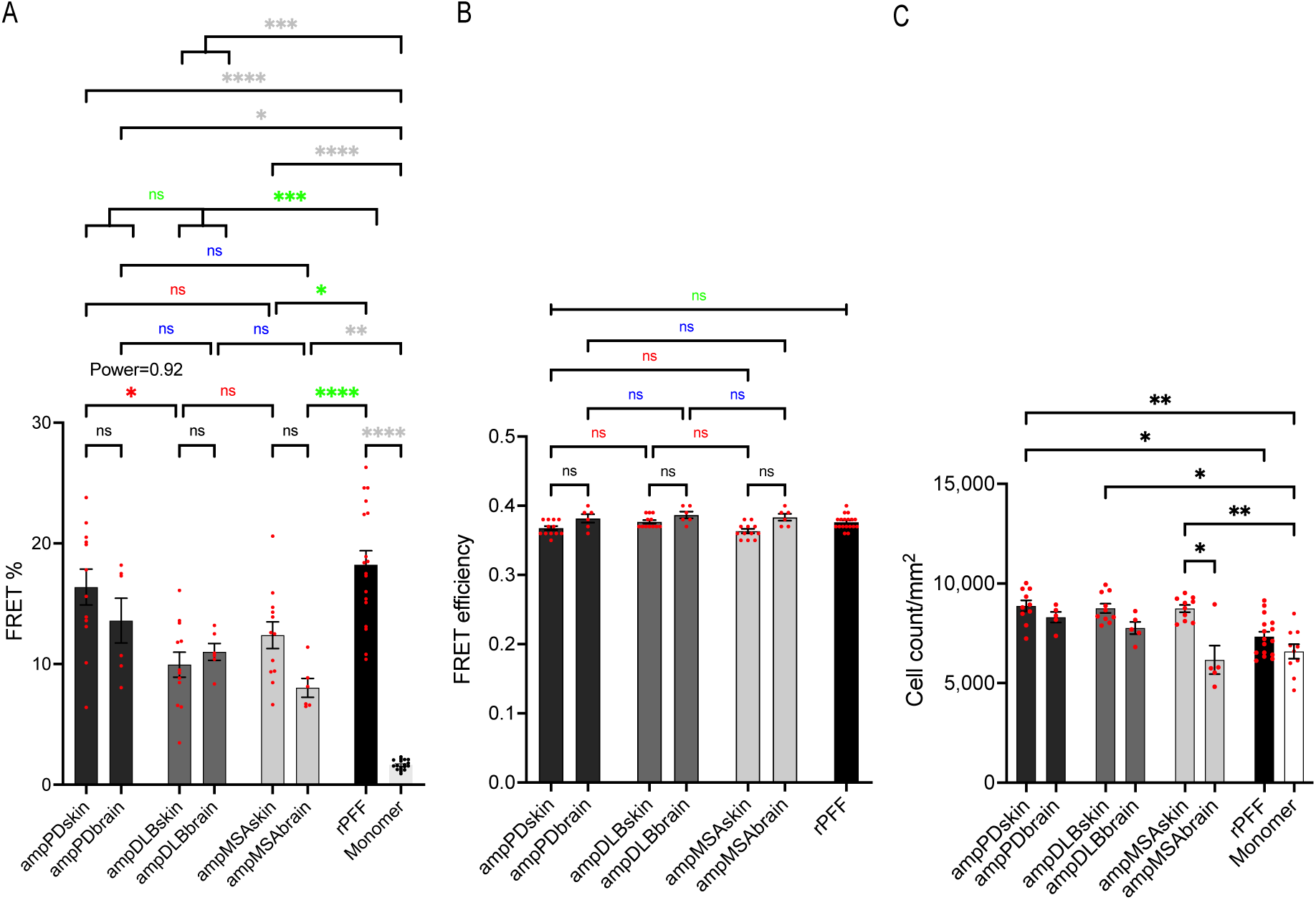
Skin- or brain-amplified strains induce intracellular αSyn aggregation triggering FRET. (**A, B, C**) Quantification of percent FRET positive cells and FRET efficiency by FRET-flow cytometry, and total cell counts for p-αSyn inclusion analysis by high-content confocal imaging. Data are presented as mean ± SEM, and each dot corresponds to data analyzed from 20,000 to 25,000 cells (A, B) in a biological replicate from 2-3 independent experiments; N=10-12 from the skin of 2 cases, N=5-6 from the brain of 1 case, N=17-18 from rPFF group, and N=18 (A, B) and 9 (C) from monomer group. Statistical analysis was performed by Brown-Forsythe and Welch ANOVA with Dunnett’s T3 multiple comparison test (A) or Kruskal-Wallis with Dunn’s multiple comparison test (B, C). A p value <0.05 was considered significant. Significant differences are indicated by *p<0.05, **p<0.01, ***p<0.001, ****p<0.0001.

